# Biomarker-driven and gene-enhanced viral therapy for malignant peripheral nerve sheath tumor

**DOI:** 10.64898/2026.10.12.758653

**Authors:** Meghna Saxena, Love Li, Brady G. Strittmatter, Kyle Williams, Faruck L. Hakkim, Shraddha Potti, Alanna Halloran, Elsie Warren, Danielle Maryanski, Courtney Barnes, Christopher Stehn, Madilyn R. Stahl, Chang-Jung Lee, Alex Larsson, Zach Seeman, Wendy Hudson, David A. Largaespada, Robert T. Galvin

## Abstract

**Background:** High-grade malignant peripheral nerve sheath tumors (MPNST) frequently lose Polycomb Repressive Complex 2 (PRC2), reducing the repressive histone mark, H3K27me3. We hypothesized that H3K27me3 loss sensitizes MPNST to oncolytic HSV due to impaired transcriptional resistance. HSV replication is also restricted by the integrated stress response, normally countered by HSV ICP34.5, representing an additional resistance pathway to *ICP34.5*-attenuated HSV.

**Methods:** PRC2 loss-of-function was evaluated as a biomarker of HSV G47Δ (*ΔICP34.5*) susceptibility in MPNST using isogenic, pharmacologic inhibition, and genetic rescue models. CUT&RUN and RNA-sequencing defined PRC2-dependent viral and host responses. We engineered a G47Δ-derived vector (G47Δ_2_) with tumor-regulated *ICP34.5* expression [G47Δ_2_,ins(*BIRC5>ICP34.5*)] and evaluated efficacy in MPNST xenografts.

**Results:** H3K27me3 deficiency increased G47Δ sensitivity and viral spread *in vitro*. CUT&RUN demonstrated PRC2-dependent H3K27me3 accumulation on the viral genome, explaining G47Δ activity in H3K27me3-deficient MPNST. RNA sequencing of G47Δ infected MPNST cells revealed heightened integrated stress response activity and blunted induction of translation-associated programs in the PRC2-deficient state (both *p*<0.001), consistent with compensatory resistance at the level of viral protein translation. Accordingly, tumor-regulated ICP34.5 HSV limited daily tumor-volume growth to 0.6% (95% CI, −2.9–4.2%), improving upon both G47Δ_2_ (8.3%; 4.5–12.2%) and controls (22.5%; 17.4–27.8%). A murine gastrointestinal toxicity phenotype reduced all-cause event-free survival, indicating vector and dosing optimizations are needed to widen the therapeutic window.

**Conclusions:** H3K27me3 loss creates a paradoxical antiviral state that tumor-selective ICP34.5 overcomes to rescue HSV anti-tumor activity, supporting biomarker-driven viral engineering to exploit the permissive transcriptional state of H3K27me3-deficient MPNSTs.

**Key Points:**

1. Loss of a key epigenetic mark (H3K27me3) MPNST is a biomarker of susceptibility to oncolytic HSV
2. H3K27me3-deficient MPNST mounts a compensatory stress response that limits HSV propagation.
3. Engineered viral gene therapy to overcome the stress response rescues antitumor activity.

**Importance of the Study:** High-grade MPNST lacks effective systemic therapies, and biomarkers that can guide oncolytic virotherapy are poorly defined. This study identifies loss of PRC2-dependent H3K27me3 as a marker of susceptibility to oncolytic HSV but shows that this same epigenetic state also primes a compensatory integrated stress response that limits productive viral propagation at the level of protein translation. These findings connect a tumor-specific chromatin defect to both enhanced viral permissiveness and a compensatory resistance mechanism. By restoring tumor-selective expression of *ICP34.5*, an HSV-encoded antagonist of the integrated stress response frequently removed to promote oncolytic vector attenuation, we demonstrate that this second barrier can be therapeutically targeted to improve antitumor activity. The work therefore provides a mechanistic framework for matching viral engineering to tumor epigenetic state. Robust anti-tumor activity must be balanced with therapeutic safety, providing avenues for further vector optimization. This biomarker-driven strategy may be relevant to other H3K27me3-deficient nervous system tumors.

## BACKGROUND

Oncolytic viruses are engineered for tumor-selective replication, with regulatory approvals of talimogene laherparepvec (T-VEC) and teserpaturev (HSV G47Δ) demonstrating the clinical feasibility of this approach.^1–4^ Their therapeutic effects include direct tumor-cell lysis, induction of antitumor immunity, and delivery of therapeutic gene cargo.^5^ Clinical studies demonstrate favorable safety, including with administration to sensitive sites like the pediatric brainstem.^3,4,6–8^ However, efficacy remains variable and durable responses uncommon. Identifying predictive biomarkers of viral sensitivity and tumor-cell resistance mechanisms remains critical for improving oncolytic virotherapy.

One compelling candidate for a biomarker of oncolytic virotherapy sensitivity is the global loss of Histone H3 Lysine 27 trimethylation (H3K27me3), a hallmark of high-grade malignant peripheral nerve sheath tumor (MPNST).^9,10^ High-grade MPNST commonly acquires loss-of-function alterations in Polycomb Repressive Complex 2 (PRC2), frequently through *SUZ12* or *EED* mutations,^11,12^ resulting in global H3K27me3 loss and epigenetic reprogramming.^13,14^ PRC2 also deposits repressive H3K27me3 on HSV DNA-associated histones during viral latency establishment.^15^ With oncolytic virotherapy, epigenetic silencing may represent a critical barrier to efficacy, as robust lytic replication is desired in this context. Therefore, H3K27me3 loss is hypothesized to enhance tumor-selective oncolytic herpesvirus efficacy via viral genome transcriptional permissiveness.

We tested this hypothesis using isogenic MPNST models with and without PRC2 activity (**Table 1**) and HSV G47Δ.^4,16^ An attenuating feature of G47Δ is deletion of *ICP34.5*, whose protein product counteracts the integrated stress response (ISR) by promoting dephosphorylation of eIF2α and rescue of protein translation during infection.^17–19^ While the latter function of ICP34.5 is desirable within cancer cells for oncolytic virotherapy efficacy, ICP34.5 also binds Beclin 1, inhibiting its autophagy function, which contributes to neurovirulence. For this reason, early generation HSV oncolytic vectors attenuate *ICP34.5*.^20^ Tumor-restricted restoration of *ICP34.5* or related ISR-antagonizing functions has therefore been pursued as a strategy to preserve oncolytic potency while limiting neurovirulence. ^21–25^ Tumor-selective promoters are frequently utilized to bias viral gene expression toward target cells. Prior studies have validated the *BIRC5 (Survivin*) promoter for this purpose, and single-cell atlases demonstrate its activity is primarily restricted to replicating malignant cells.^26–32^

**Table 1:**
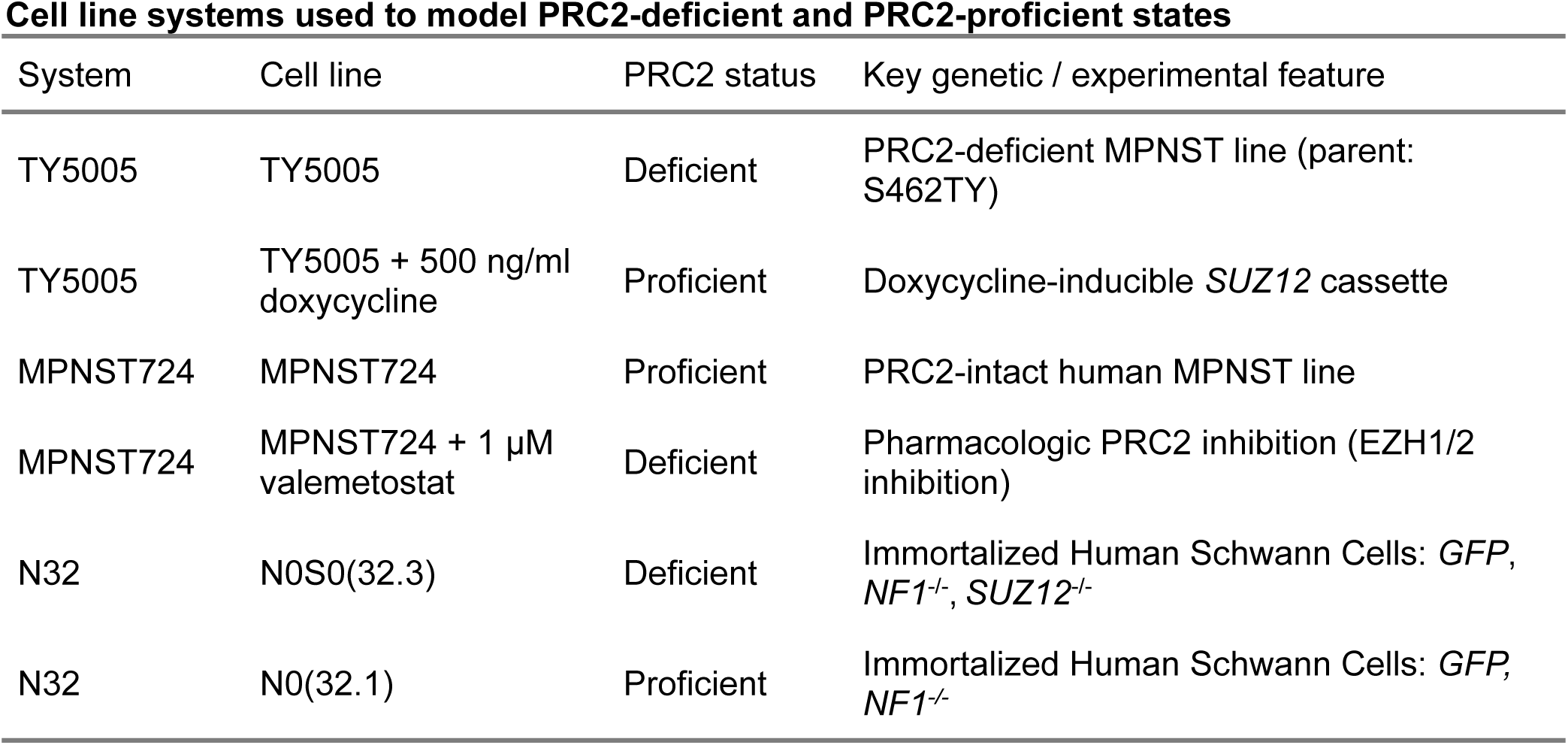
Three cellular systems were used to model paired PRC2-proficient and PRC2-deficient states in MPNST and MPNST-relevant Schwann-cell models, including inducible genetic rescue, pharmacologic inhibition, and isogenic genetic deletion models. For the inducible rescue system, S462-TY parental cell line contains a doxycycline inducible TET cassette (TY5005).^33^ TY5005 cells were pre-treated with doxycycline (500 ng/ml) for a minimum of 14 days to restore *SUZ12* expression and PRC2 activity. For the pharmacologic inhibition system, MPNST724 cells (PRC2 proficient) were pre-treated with the EZH1/2 inhibitor valemetostat (1 µM) for a minimum of 7 days to induce PRC2 loss-of-function. N0(32.1) represents *hTERT*-immortalized human Schwann cells with *NF1* deletion via CRISPR Cas9. N0S0(32.3) is an isogenic derivative of N0(32.1) characterized by additional *SUZ12* knock-out.^13^

Here, we identify a paradoxical antiviral state in H3K27me3-deficient MPNST. Loss of PRC2 reduces epigenetic restriction of the HSV genome and enhances viral permissiveness at the transcriptional level. However, PRC2 loss is also associated with a compensatory increase in the integrated stress response, highlighting a translational resistance mechanism that is consequential in the H3K27me3-deficient epigenetic state. Engineering tumor-selective *ICP34.5* expression overcomes this second barrier and markedly enhances antitumor activity in H3K27me3-deficient MPNST. Together, these findings provide a biomarker-driven strategy to exploit defective viral genome silencing while circumventing compensatory innate translational resistance.

## METHODS

### Cell culture and maintenance

Cells were cultured in DMEM with 10% FBS and 1% penicillin/streptomycin at 37°C and 5% CO₂. Cultures were routinely confirmed mycoplasma-negative (MycoAlert, Lonza). Primary human astrocytes (iXCell, #10HU-188, Lot 204049) were cultured in Schwann cell growth medium (iXcell, #MD-0055) on flasks coated with hESC-Qualified matrigel (Corning, #08-774-552).

### Cell line engineering

TY5005 is nomenclature for the S462-TY parental cell line containing a doxycycline inducible TET cassette as previously described.^33^ N0(32.1) (*NF1^-/-^; GFP^+^*) and N0S0(32.3) (*NF1^-/-^; SUZ12^-/-^; GFP^+^*) were generated from *hTERT*-immortalized human Schwann cells as previously described.^13^ A *Suz12*-knock-out derivative of the NP^cis^ C57BL/6 syngeneic MPNST model is described in Supplementary methods.

### Oncolytic HSV amplification, purification, and titration

HSV G47Δ, HSV-1 F (ATCC), and engineered G47Δ derivatives were amplified in Vero cells, purified by sucrose-gradient centrifugation, and titrated by plaque assay according to the protocol published by Nguyen et al.^34^ G47Δ-derived viruses expressing β-galactosidase were visualized with X-Gal, whereas HSV-1 F plaques were stained with Giemsa. Engineered viral Bacterial Artificial Chromosomes (BACs) were stored in DH10B E. coli glycerol stocks, amplified in LB media, and purified using the NucleoBond Xtra Midi kit (Macherey-Nagel). Sequence verified purified plasmid (PlasmidSaurus) was transfected into Vero cells by 4D-Nucleofection (Lonza; SF solution and Supplement; 400,000 cells and 1 µg DNA per reaction, and DN-100 pulse code). After visualization of plaque formation, recovered virus was subsequently amplified, purified, and titrated as above.

### Cell viability assays

Cells were seeded at 3,000–5,000 cells/well in 96-well plates and infected the following day with serially diluted HSV inocula. Experiments comprised three independent biological replicates, each with six technical replicate wells per multiplicity of infection (MOI). After 1.5hr adsorption, inoculum was replaced with complete medium. Viability was measured 6 days after infection using CellTiter 96 AQueous One Solution (Promega). Absorbance was measured at 450 and 690 nm using a Cytation 5 reader (BioTek), background corrected and normalized to matched, uninfected controls.

### Flow cytometry

Viral gene expression was assessed using empirically optimized HSV MOIs. At specified time points, cells were stained with Fixable Viability Dye eFluor 780 and Spider-βGal (Dojindo), fixed in 2% formaldehyde, and acquired on a CytoFLEX S (Beckman Coulter). Uninfected and dead-cell controls were included to define β-galactosidase and viability gates. Data were analyzed in FlowJo v10.10.0 after exclusion of debris and doublets.

### Western Blotting

Cells were lysed in RIPA buffer supplemented with protease and phosphatase inhibitors, and protein concentration was determined by BCA assay. Equal amounts of lysate were resolved by SDS-PAGE and transferred to PVDF membranes. Membranes were blocked in TBS-T containing 5% non-fat dry milk or 5% BSA for phosphorylated targets and incubated overnight at 4 °C with primary antibodies, followed by HRP-conjugated secondary antibodies. Bands were detected using WesternBright Quantum chemiluminescent substrate (Advansta) and imaged on a LI-COR system.

Antibodies from Cell Signaling Technology included: phospho-eIF2α (rabbit mAb #9721; 1:1000), total eIF2α (rabbit mAb #9722; 1:1000), H3K27me3 (rabbit mAb #9733; 1:1000), GAPDH (rabbit mAb #5174; 1:1000), β-actin (rabbit mAb #4967; 1:1000), and Vinculin (rabbit mAb #13901; 1:1000). Secondary antibody used was HRP-linked anti-rabbit IgG (#7074; 1:5000).

### CUT&RUN

TY5005 cells were used to assess PRC2-dependent occupancy of H3K27me3, H3K4me3, and GST-MeCP2 on host and viral genomes. Biological replicates (n=3 per PRC2-rescue × infection condition) were infected with HSV G47Δ at an MOI of 0.1. At 24 hours, viability and viral gene expression were assessed by flow cytometry (**Supp. Fig. 2**), and nuclei were isolated using CUTANA Extraction Buffer supplemented with 0.5 mM spermidine and protease inhibitors (EpiCypher #14-1048) before cryopreservation at −80 °C.

CUT&RUN Assay and Sequencing: CUT&RUN was performed by EpiCypher Services using the automated CUTANA ChIC/CUT&RUN workflow with 500,000 nuclei per reaction (48 reactions).^35^ Concanavalin-A-immobilized nuclei (EpiCypher #21-1401) were incubated overnight at 4 °C with antibodies against H3K27me3 (EpiCypher #13-0055), H3K4me3 (EpiCypher #13-0060), or matched IgG control (EpiCypher #13-0042); DNA methylation was profiled using recombinant GST-MeCP2 (EpiCypher #15-2002) followed by anti-GST targeting (EpiCypher #13-0073).^36^ Chromatin cleavage was performed with pAG-MNase (EpiCypher #15-1016), with 0.5 ng *E. coli* DNA (EpiCypher #18-1401k) per 500,000 nuclei added for spike-in normalization. The SNAP-CUTANA K-MetStat panel of barcoded nucleosomes carrying defined methylation states was included to monitor assay specificity. Released DNA was bead-purified, normalized to 5 ng when sufficient material was recovered, prepared using the CUTANA CUT&RUN Library Prep Kit (EpiCypher #14-1001), and sequenced on an Illumina NextSeq 2000 (2×50 bp paired-end reads).

Reads were aligned with Bowtie2 to a concatenated GRCh38, HSV-1 F (GU734771.1), and *E. coli* K-12 (GCF_000005845.2) reference and filtered at MAPQ ≥30 using SAMtools. Coverage was normalized using a sample-specific *E. coli* scaling factor (100,000/*E. coli*-aligned reads), and 10-bp BigWig tracks were generated with deepTools. Because infected samples showed substantial nonspecific HSV signal in matched IgG controls, normalized IgG signal was subtracted from H3K27me3, H3K4me3, and GST-MeCP2 tracks using bigwigCompare. IgG-subtracted signal was quantified using multiBigwigSummary in 10-kb bins across HSV-1 and 100-kb bins across the human genome; representative tracks were visualized with pyGenomeTracks. Assay-specific technical QC failures were excluded as detailed in Supp. Fig. 2.

### RNA sequencing and transcriptomic analysis

RNA-sequencing was performed across paired PRC2-proficient and PRC2-deficient states of the TY5005, MPNST724, and N32 systems, with six independent biological replicates per PRC2-status and infection condition (N=72 samples). Replicates were cultured independently and infected with empirically optimized, sublethal HSV G47Δ doses. Because PRC2 status altered HSV replication kinetics, samples were harvested using a viral-burden–matched design. Beginning 24hr after infection, aliquots from the six biological replicates within each condition were pooled solely for viral-burden assessment (**Supp. Fig. 3**). Samples were harvested once the pooled infected population reached >80% viral β-galactosidase positivity while maintaining viability, with matched uninfected controls collected at the corresponding time point. Collection occurred 36–72hr after infection depending on the model and PRC2 state. Viral spread was assessed using Spider-βGal in TY5005 and MPNST724 cells. In N32 cells, constitutive GFP expression interfered with this assay, so infection was estimated semi-quantitatively by X-gal staining. RNA was isolated using the RNeasy Kit (Qiagen). RNA quality was assessed by RiboGreen fluorimetry and capillary electrophoresis; samples with >500 ng RNA and RIN ≥8 underwent stranded mRNA library preparation using the Watchmaker mRNA Library Preparation Kit with oligo-dT enrichment. Libraries were sequenced by the University of Minnesota Genomics Center across three AVITI Freestyle High 2 × 150-bp runs.

Raw FASTQ files were quality assessed using FastQC/MultiQC, trimmed with fastp, and aligned STAR v2.7.11b to a combined human GRCh38/GENCODE v46 and HSV-1 F (GU734771.1) reference. Gene-level counts were generated using featureCounts (Subread v2.1.1) for reversely stranded paired-end reads. Viral genes were excluded from the host DESeq2 matrix and analyzed separately to prevent abundant HSV transcripts from distorting host-gene size-factor normalization. Gene sets were retrieved programmatically from Hallmark, Reactome, and Gene Ontology (GO) Biological Process collections in MSigDB using msigdbr. Sample-level pathway activity was quantified by single-sample Gene Set Enrichment Analysis (ssGSEA) using GSVA. Integrated stress response burden was additionally summarized as the mean variance-stabilized (VST) expression of GO ISR genes, and a global Translation/RNA activity index was calculated as the mean z-standardized ssGSEA score across MSigDB pathways matching translation, ribosome biogenesis, and RNA-processing annotations containing 15 – 500 genes.

### Dual Luciferase Assay

Cells were co-transfected using Lipofectamine 3000 with firefly luciferase (Luc2) reporter plasmids driven by the minimal promoter (*MinP*), *CBh* promoter, *BIRC5* promoter, or *NES* enhancer element, and an SV40-driven pRL *Renilla* luciferase control vector (Promega) at a 50:1 firefly:Renilla plasmid ratio. At 24 hours after transfection, firefly and Renilla luciferase activities were measured using the Dual-Luciferase Reporter Assay System (Promega). Firefly luminescence was normalized to Renilla luminescence for each sample, and promoter activity was expressed relative to the CBh>Luc2 positive-control construct, defined as 100%.

### Bacterial Artificial Chromosome Recombineering

An HSV G47Δ BAC (HSV G47Δ_2_) was first generated from the BeloBAC-HSV(1)-F BAC by recombineering.^16^ Subsequently, a reverse-oriented cassette (SV40pA:pBIRC5>ICP34.5:rBGpA-CBh>tTS:SV40pA) was inserted into the two deleted *ICP34.5* loci to generate HSV G47Δ_2_,ins(BIRC5>ICP34.5). The CBh>tTS element is functionally inert in the context of this study but leveraged in separate, unpublished derivatives. Full vector construction protocols are detailed in the Supplementary Methods. Briefly, targeted genomic modifications were achieved via homologous recombination in *E. coli* utilizing established selection and counter-selection strategies.^37,38^

### Polymerase Chain Reaction (PCR) and agarose gel electrophoresis

DNA was purified using the QIAamp DNA Mini Kit (QIAGEN) according to the manufacturer’s protocol. PCR was performed using AccuPrime Taq DNA Polymerase, High Fidelity or AccuPrime GC-Rich DNA Polymerase (Thermo Fisher Scientific), selected according to template characteristics, with cycling conditions optimized for individual primer pairs. Amplicons were resolved on ethidium bromide-containing agarose gels and imaged using a ChemiDoc XRS+ system (Bio-Rad).

### In vivo experiments

To establish subcutaneous xenografts, athymic nude mice were flank-injected with 2.0×10⁶ S462-TY cells in 200 µL 1:1 PBS:Matrigel. Following implantation, mice were pre-randomized in R into respective study arms (N = 10 control, N = 12 per viral group). Pre-defined endpoints were tumor volume of 1000 mm^3^, and mice survival was to be assessed at day +30 from enrollment. Confirmed response was defined as a ≥20% decrease from the preceding observed tumor maximum sustained at the next measurement. Pilot studies suggested enhanced activity from a multi-dosing strategy, and the chosen dose (1.0×10^7^ PFU/dose) was chosen to approximate the intratumoral dosing of rQNestin34.5 in CNS preclinical toxicology evaluations.^39^

Enrollment occurred once tumors reached a volume of 100-125 mm³, calculated using the ellipsoid formula: 0.5 × (width² × length). Mice were treated on Day 1 (enrollment) and Day 4 with 50 µL of intratumoral injection of either PBS or respective virus. Tumor dimensions were measured daily by investigators blinded to the treatment group assignments. Animal weights were recorded every two days beginning on the day of enrollment. During the trial, unexpected gastrointestinal toxicity (ileus and toxic megacolon) was observed in a subset of virus-treated mice. To mitigate this toxicity proactively, the drinking water of remaining mice was supplemented with polyethylene glycol 3350 (Miralax) per Khoury-Hanold, et al.^40^ The initial concentration of 15% w/v was subsequently reduced to 7.5% w/v due to secondary weight loss and dehydration. Pre-defined euthanasia criteria included a tumor volume exceeding 1000 mm³ or a reduction in body weight of ≥20% from baseline.

Upon euthanasia, all available tumor and liver tissues (N = 4 control samples were randomly selected) were harvested for immunohistochemical (IHC) analysis. All staining procedures performed at HistoWiz, Inc, using the Leica Bond RX automated stainer (Leica Microsystems). Samples were processed, embedded in paraffin, and sectioned at 4 μm. Slides were dewaxed using xylene and alcohol based dewaxing solutions. Epitope retrieval was performed by heat-induced epitope retrieval of the formalin-fixed, paraffin-embedded tissue. Endogenous peroxidase was blocked using peroxide block buffer. Tissue was incubated with the H3K27me3 (Ozyme #CST9733), β-Galactosidase (ThermoFisher Scientific #14-6773-81), Ki67 (Ozyme #CST34330) antibodies, followed by DAB rabbit secondary reagents according to the manufacturer’s protocol. Whole-slide images were downloaded as SVS files and quantitatively analyzed using QuPath.

### Statistical Analysis

Statistical analyses were conducted using R version 4.4.1 and GraphPad Prism v10. For *in vitro* viability assays, dose-response curves were fitted by nonlinear regression and IC_50_ values compared using the extra sum-of-squares F test. Other comparisons used Welch’s *t*-tests or Kruskal–Wallis tests with Dunn’s post hoc comparisons, with BH or Holm correction for multiple testing as appropriate. Statistical tests were two-sided.

For RNA-sequencing analyses, genes with fewer than 100 total counts across samples were excluded. Host differential expression was modeled using DESeq2 with the design ∼ System + PRC2_status + infection + PRC2_status:infection; viral transcripts were analyzed separately among infected samples. Differentially expressed genes were defined by FDR <0.05 and |log2 fold-change| ≥1. Genes were ranked by the DESeq2 test statistic for fast preranked gene set enrichment analysis (FGSEA). GO Integrated Stress Response, Hallmark Interferon-α Response, and Hallmark Apoptosis were selected for focused pathway analysis based on the study hypothesis and prior literature. FGSEA provided normalized enrichment scores, while significance for focused between-phenotype comparisons was assessed using BH-adjusted q-values from corresponding system-adjusted ssGSEA models with HC3 robust covariance estimates. Sample-level ssGSEA scores, the VST-derived ISR score, and the global Translation/RNA activity index were analyzed using factorial linear models incorporating System, PRC2 status, infection, and their interaction, with HC3 heteroscedasticity-robust covariance estimates (sandwich, lmtest). Estimated marginal means and prespecified contrasts were obtained using emmeans. For the VST-derived ISR analysis, partial η² and Cohen’s *d* were reported as effect-size measures for model terms and covariate-adjusted contrasts, respectively. Viral differential expressions were analyzed independently among infected samples using a system-adjusted model (∼ System + PRC2_status). CUT&RUN spike-in-normalized, IgG-subtracted genomic occupancy was analyzed using Type II two-way factorial ANOVA (Signal ∼ PRC2 rescue × Infection), with post-hoc contrasts estimated using emmeans.

For *in vivo* analyses, longitudinal tumor growth was evaluated using a linear mixed-effects model of log1p-transformed tumor volume (log(Volume+1) ∼ Day × Treatment + Gender + (Day | MouseID)), incorporating mouse-specific random intercepts and slopes. Treatment-specific growth slopes were estimated using emtrends, with Tukey-adjusted pairwise comparisons. All-cause event-free survival, in which tumor endpoint or toxicity-related euthanasia constituted an event, was estimated by the Kaplan-Meier method and compared using an overall log-rank test and BH-adjusted pairwise log-rank tests. Cumulative-incidence functions for tumor endpoint were estimated in the presence of competing events and compared among treatment groups using Gray’s test. Body-weight change was calculated relative to baseline and summarized by treatment and study day as mean ± SD. Exploratory delayed-response kinetics within the BIRC5-regulated ICP34.5 treatment group were evaluated using piecewise linear mixed-effects models of log1p-transformed tumor volume. Candidate change points were compared by maximum-likelihood model fitting, with the minimum-AIC breakpoint selected and an approximate 95% profile-likelihood support interval calculated.

## RESULTS

### Loss of H3K27me3 sensitizes MPNST to oncolytic HSV infection and accelerates viral spread

All evaluated human MPNST cell lines were sensitive to HSV G47Δ (IC_50_ <0.1 MOI), exhibiting significantly greater sensitivity than *hTERT*-immortalized human Schwann cells (**Fig. 1A**). We next evaluated the effect of H3K27me3 status using paired genetic, pharmacologic, and isogenic model systems (**Table 1**). *SUZ12* loss in human *hTERT*-immortalized Schwann cells shifted HSV G47Δ dose-response curves leftward (*p*<0.001; **Fig. 1B**). A *Suz12*-deleted derivative of the syngeneic NPcis MPNST line JW18 (JW18_S6) demonstrated similar increased viral sensitivity, although this murine model was overall less permissive to HSV than human models (**Supp. Fig. 1**). In PRC2-proficient MPNST724 cells, pharmacologic H3K27me3 depletion with valemetostat significantly decreased the IC_50_ (*p*<0.001), but did not alter sensitivity of H3K27me3-deficient S462-TY (*p*=0.8; **Fig. 1C**). Conversely, doxycycline-induced SUZ12 restoration and H3K27me3 recovery in TY5005 cells shifted the dose-response curve rightward (*p*<0.001; **Fig. 1D**).

**Figure 1:**
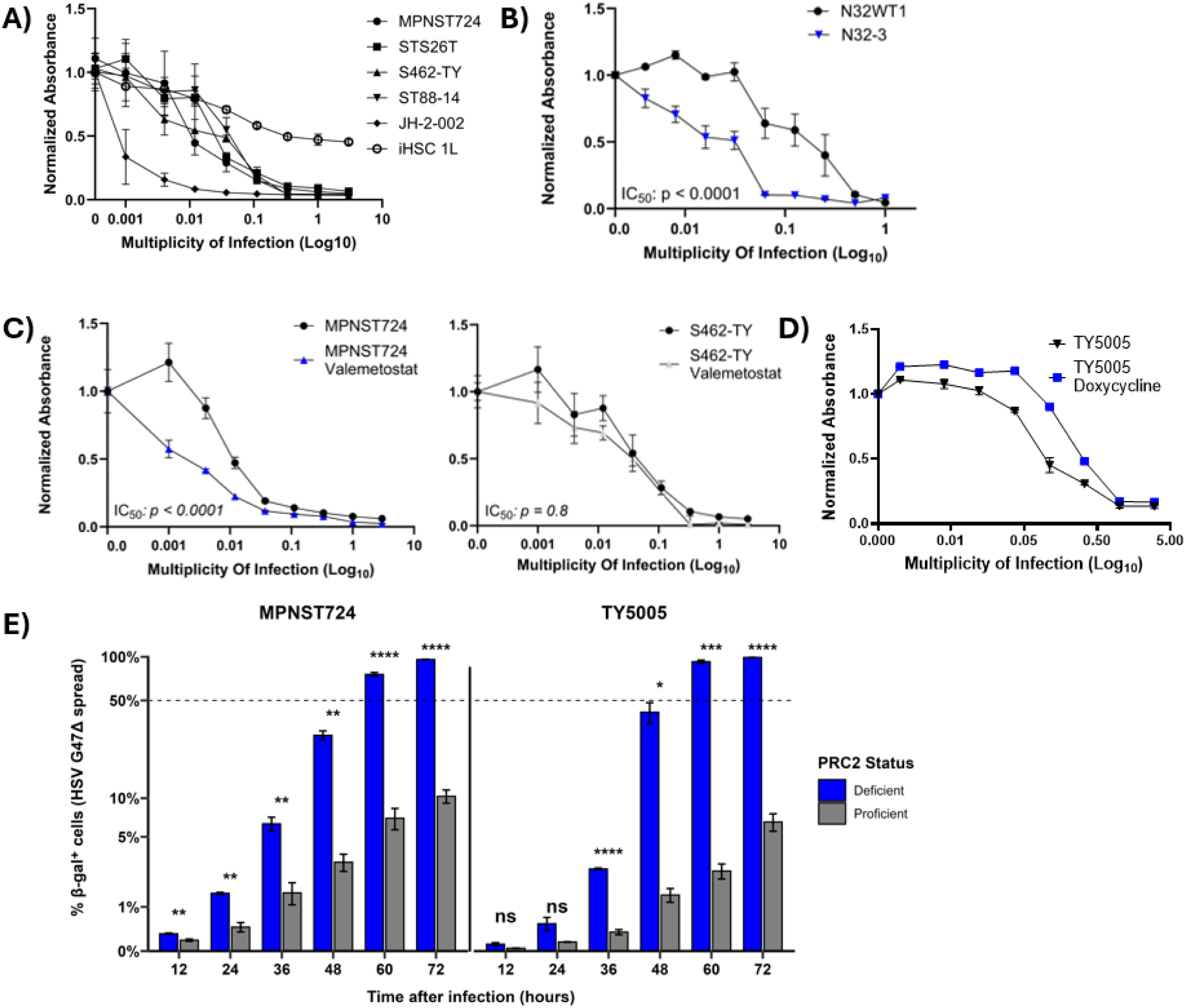
MPNST is permissive to oncolytic herpes simplex virus, and loss of H3K27me3 further sensitizes MPNST to productive lytic infection. **A-D)** MTS cell viability assays are presented with viability day +6 from infection normalized to uninfected controls. Error bars represent standard error of the mean for biological replicates, where each experiment utilized N = 6 technical replicates per dose. Blue lines and bars correspond to the H3K27me3-deficient condition. Dose-response curves were fitted via non-linear regression, and IC_50_ values were compared using the Extra sum-of-squares F test. A) HSV G47Δ demonstrates significantly greater cytotoxicity against all evaluated human MPNST cell lines compared to *hTert*-immortalized primary human Schwann cells (iHSC 1L), based on IC_50_ comparisons (all *p*<0.001). The mean IC_50_ for every tested MPNST cell line is <0.1 MOI. B) In isogenic models of MPNST derived from *hTert*-immortalized human Schwann cells, *SUZ12* knock-out shifts the IC_50_ to the left (*p*<0.001). C) PRC2-proficient MPNST724 is further sensitized to HSV G47Δ when H3K27me3 is depleted by pharmacologic pretreatment with valemetostat (*p*<0.001), while drug pretreatment did not significantly alter the IC_50_ in PRC2-deficient S462-TY (*p*=0.8). D) S462-TY was engineered to restore *SUZ12* under a doxycycline-inducible TET system (TY5005 is the derivative of parental S462-TY). After genetic rescue restoring H3K27me3, the IC_50_ of HSV G47Δ significantly shifted rightward (*p*<0.001). E) Equally seeded MPNST724 (valemetostat pretreated vs. wildtype) and TY5005 (doxycycline induced vs. wildtype) were infected at 0.01 MOI. Adherent cells were collected at indicated time points, and the percentage of β-galactosidase positive cells was quantified by flow cytometry. Kinetics of HSV G47Δ spread were significantly increased in the H3K27me3-deficient context. By 72 hours, 100% infection coverage was observed in the H3K27me3-deficient condition compared to approximately 10% in the PRC2-proficient condition. Error bars represent SEM for three biological replicates. Statistical significance between PRC2-proficient and deficient conditions at each timepoint was determined using Welch’s t-tests, with p-values adjusted for multiple comparisons (BH procedure) [*: *p*≤0.05; **: *p*≤0.01; ***: *p*≤0.001; ****: *p*≤0.0001].

H3K27me3 status also altered viral spread. Following 0.01 MOI infection, β-galactosidase-positive cells accumulated much more rapidly in PRC2-deficient MPNST724 and TY5005 conditions, with significant differences evident by 36hr. By 72hr, ∼100% of cells were β-galactosidase-positive in PRC2-deficient conditions compared with ∼10% in matched PRC2-proficient conditions (**Fig. 1E**).

### PRC2 restoration increases H3K27me3 occupancy on the HSV-1 genome, highlighting an impaired resistance mechanism in H3K27me3-deficient MPNST

Because PRC2 can establish repressive chromatin, loss of function could represent a therapeutic vulnerability (**Fig. 2A**). Immunoblotting demonstrated increased total cellular H3K27me3 following HSV infection in PRC2-proficient conditions by 12 hours post-infection (**Fig. 2B**). We asked whether infection-associated increases in H3K27me3 reflect deposition on host or viral genomes. In the TY5005 CUT&RUN model, doxycycline-mediated PRC2 rescue restored global H3K27me3 occupancy across the human genome, without a significant difference between infected and uninfected cells (**Fig. 2C**). In contrast, PRC2 rescue produced significant H3K27me3 occupancy across the HSV-1 genome following infection (**Fig. 2D**), distributed broadly across mappable viral sequences (**Fig. 2E**). These findings support PRC2-dependent HSV DNA chromatinization as a potential barrier to productive infection. Exploratory H3K4me3 and GST-meCP2 profiling demonstrated additional differences in host- and viral-genome occupancy following infection (**Supp. Fig. 2**).

**Figure 2:**
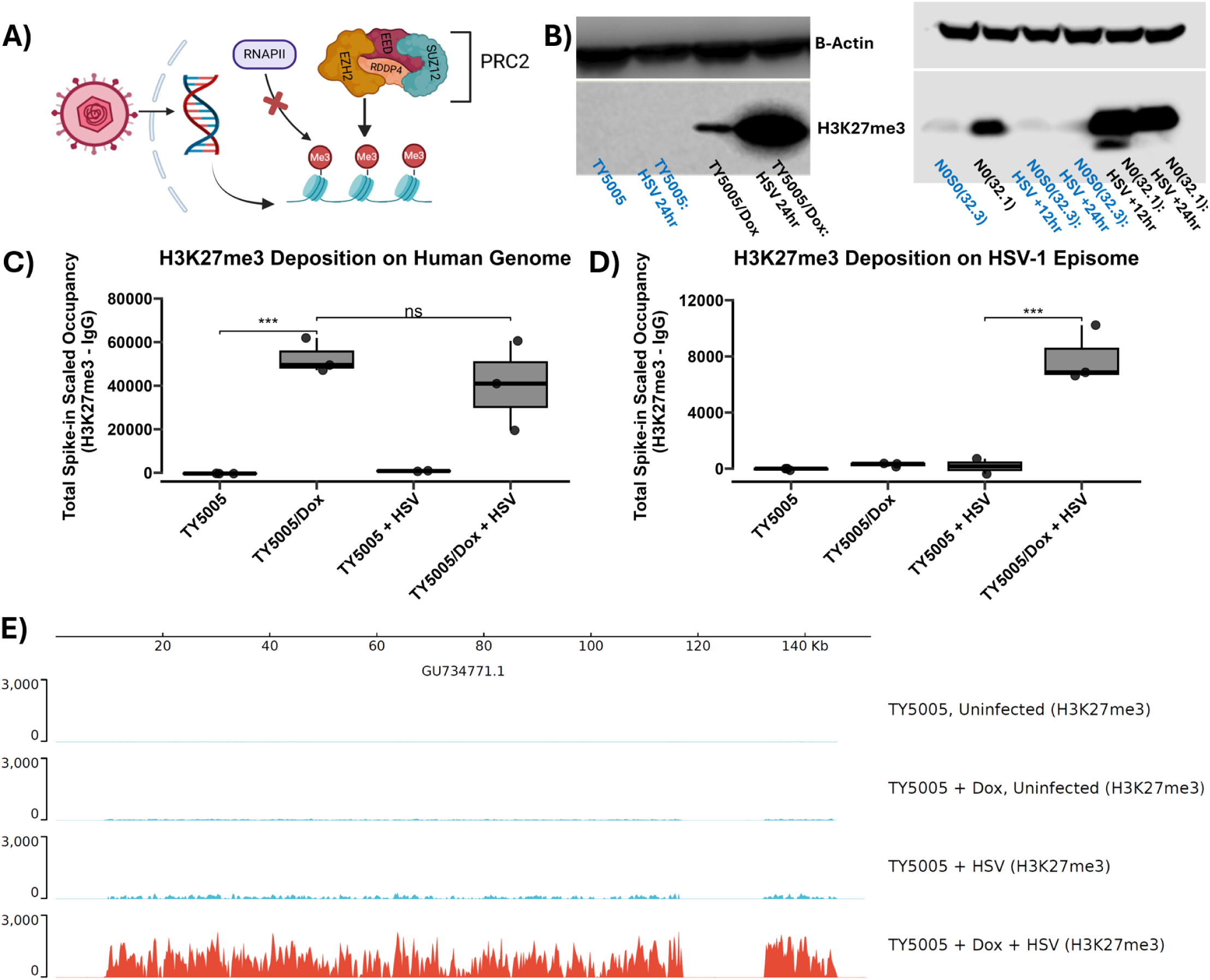
H3K27me3-deficient MPNST has intrinsic sensitivity to oncolytic virotherapy secondary to transcriptional permissiveness. **A)** Schematic representation of the proposed viral silencing mechanism, wherein the host Polycomb Repressive Complex 2 (PRC2) deposits H3K27me3 on host histones associated with viral DNA, reducing RNA polymerase II (RNAPII) transcription. **B)** Excess total cell H3K27me3 immunoblotting is observed in doxycycline-induced TY5005 (TY/Dox) at 24 hours post-infection (left). Similarly, excess H3K27me3 is observed in the infected PRC2-proficient N0(32.1) at both 12 and 24 hours post-infection (right). **C-D)** CUT&RUN analysis of H3K27me3 occupancy on the human genome (C) and HSV-1 episome (D) in TY5005 with or without *SUZ12* genetic rescue. Y-axes represent total H3K27me3 genomic occupancy, normalized to an *E. coli* spike-in control and mathematically background-subtracted against matched IgG negative controls (H3K27me3 - IgG). Statistical significance was determined using a Two-Way ANOVA followed by post-hoc pairwise comparisons via estimated marginal means [***: *p*≤0.001; ns: not significant]. **C)** Quantification of H3K27me3 on the human genome confirms successful doxycycline induction of global H3K27me3, with no significant difference observed in host genome deposition between uninfected and infected TY5005/Dox biological replicates. **D)** H3K27me3 accumulation on the HSV-1 genome was detected specifically after PRC2 rescue, consistent with PRC2-dependent chromatinization of viral DNA. **E)** Representative track plot visualizing H3K27me3 occupancy across the HSV-1 genome (GU734771.1) for the conditions evaluated in panel D. Robust, widespread H3K27me3 deposition is visible across the viral episome strictly in the TY5005 + Dox + HSV condition. Regions lacking signal correspond to HSV-1 repeat regions that are excluded in multimapping alignment filtering.

### PRC2 deficiency is associated with enhanced antiviral Integrated Stress Response

Despite enhanced HSV permissiveness in vitro, pilot experiments with S462-TY xenografts failed to regress. We therefore evaluated whether PRC2 status was associated with differential activation of host antiviral stress responses (**Fig. 3A**). HSV infection induced eIF2α phosphorylation by 12hr, confirming an intact eIF2α stress response in MPNST cells (**Fig. 3B**). We next assessed the interferon alpha (IFN-α) pathway, which produces interferon stimulated genes in response to infection;^41^ the ISR effector pathway; and the downstream apoptosis pathway by performing RNA-sequencing across all three paired PRC2 systems using a viral-burden–matched design (n = 6 biological replicates per condition). Infected samples were collected after >80% viral spread, with matched uninfected controls harvested at corresponding time points (36–72hr; **Supp. Fig. 3**), to minimize confounding by differential HSV replication kinetics.

**Figure 3:**
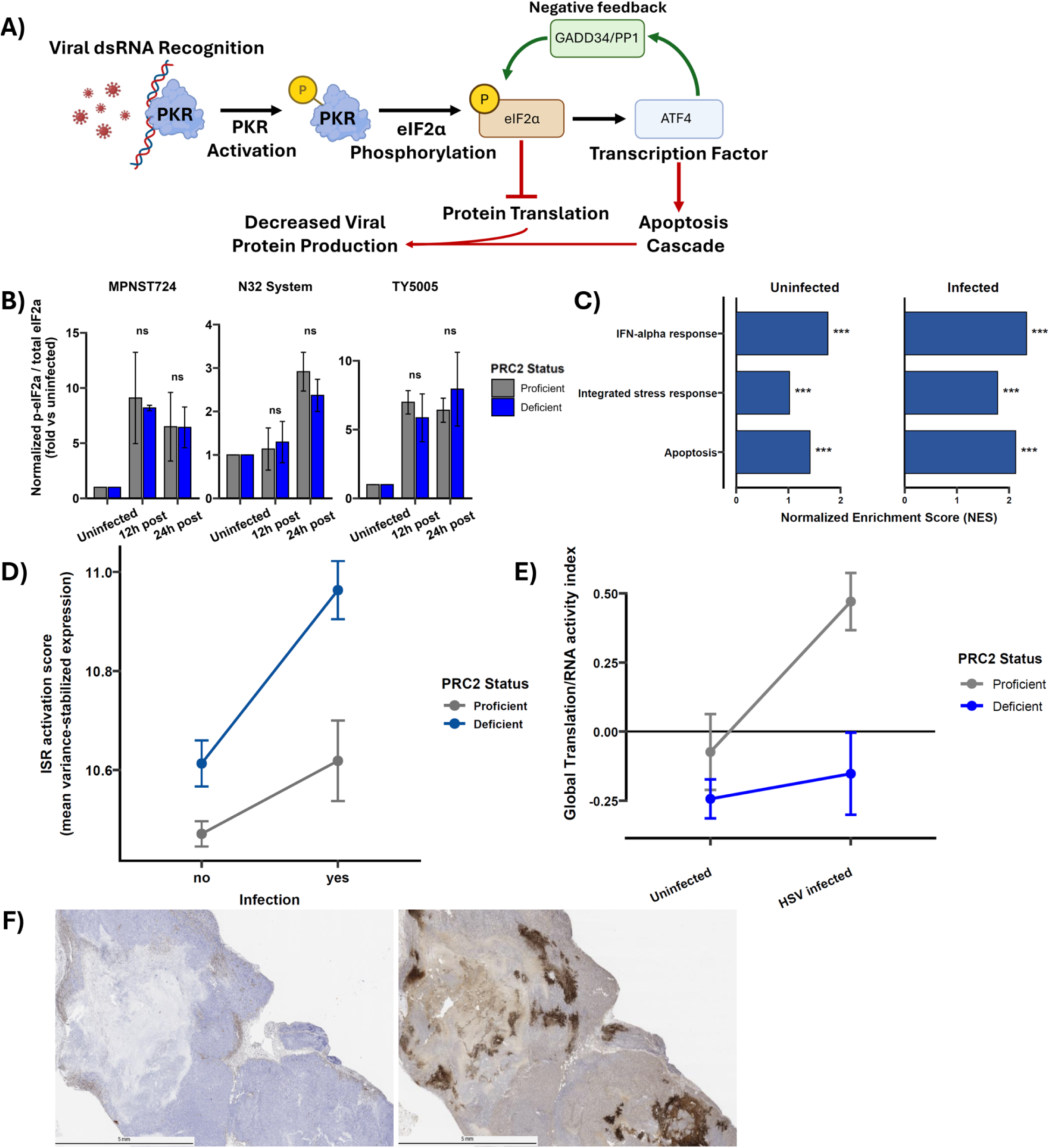
The integrated stress response represents a host innate resistance mechanism to viral infection and is hyperactivated in H3K27me3-deficient MPNST. **A)** Schematic representation of the integrated stress response. Viral double-stranded RNA (dsRNA) can activate PKR, leading to phosphorylation of eIF2α and suppression of global translation initiation. ATF4 is preferentially translated during eIF2α phosphorylation and promotes downstream apoptotic transcriptional programs, while recovery is mediated by GADD34-mediated restoration of eIF2α activity. **B)** Ratio of phosphorylated to total eIF2α in each PRC2 modular system following HSV infection, determined by immunoblotting in *n*=2 independent biological replicates per indicated time point and normalized to matched uninfected controls. Error bars indicate the range between independent experiments. HSV infection increased p-eIF2α:eIF2α across the evaluated models, consistent with activation of the eIF2α stress-response axis. No significant differences were detected between H3K27me3-deficient and proficient states at the evaluated time points (Welch’s two-sample *t*-test; ns), although this study was not powered to detect subtle differences**. C–E)** Transcriptomic responses to HSV were evaluated across the three PRC2 modular systems (n=6 biological replicates per PRC2-status and infection condition). Because viral replication kinetics differed by PRC2 phenotype, infected samples were harvested at a matched stage of viral spread (>80% β-galactosidase-positive cells), with corresponding time-matched uninfected controls. **C)** Focused FGSEA of IFN-α response, ISR, and apoptosis pathways in PRC2-deficient versus proficient cells within the uninfected and infected states. Positive normalized enrichment scores indicate enrichment in the PRC2-deficient phenotype. Bars represent FGSEA normalized enrichment scores; significance annotations reflect BH-adjusted *q-*values from the corresponding system-adjusted sample-level ssGSEA models using HC3 robust covariance estimates [***, *q*<0.001]. PRC2-deficient cells demonstrate elevated baseline activity of these antiviral/stress-response programs that persists during productive HSV infection. **D)** Sample-level ISR score, calculated as the mean variance-stabilized expression of genes within the GO ISR pathway. Estimates were obtained from a factorial linear model adjusting for model system and incorporating PRC2 status, infection, and their interaction with HC3 robust covariance estimates. Among uninfected cells, PRC2 deficiency was associated with a +0.14 VST-unit increase in ISR score (*p*<0.001; partial η²=0.55). Among PRC2-proficient cells, HSV infection increased the ISR score by +0.15 VST units (*p*<0.001; partial η²=0.56). A positive PRC2 × infection interaction (+0.20 VST, *p*<0.001; partial η² = 0.17) indicated a greater infection-associated ISR increase in PRC2-deficient cells. The adjusted contrast between proficient/uninfected and deficient/infected states was +0.49 VST units (*p*<0.001; Cohen’s *d* = 4.27). **E)** Global Translation/RNA activity index, representing the mean of z-standardized ssGSEA scores across MSigDB pathways associated with translation, ribosome biogenesis, and RNA processing. Estimates were obtained from a factorial linear model with HC3 robust covariance. HSV infection increased the index in PRC2-proficient cells (+0.54 units, *p*<0.001) but not significantly in PRC2-deficient cells (+0.09 units, *p*=0.27). The PRC2 × infection interaction was negative (−0.45, *p*<0.001), consistent with blunted infection-associated transcriptional induction of translation/RNA-associated programs in the PRC2-deficient state. **F)** Representative immunohistochemistry of an HSV G47Δ-treated S462-TY xenograft demonstrating limited viral distribution. β-galactosidase staining (brown) was concentrated near the central necrotic region, with patchy signal throughout the remaining viable tumor. Scale bar = 5 mm.

Focused pathway analysis demonstrated greater IFN-α response, ISR, and apoptosis-associated activity in PRC2-deficient cells both before and after HSV infection (**Fig. 3C**). Fast gene set enrichment analysis (FGSEA), which tests whether genes within a predefined pathway are preferentially represented among genes associated with one phenotype, yielded positive normalized enrichment scores for the PRC2-deficient state. To confirm individual-sample findings across model systems, we compared PRC2 phenotypes using system-adjusted single-sample GSEA (ssGSEA) pathway scores. PRC2-deficient cells showed significantly higher activity of all three pathways in both uninfected and infected states (BH-adjusted *q*<0.001 for all comparisons). The VST-derived GO ISR score was similarly elevated in uninfected PRC2-deficient cells (+0.14 VST units, *p*<0.001; partial η²=0.55), with a positive PRC2 × infection interaction (+0.20 VST units, *p*<0.001; partial η²=0.17), indicating greater infection-associated ISR induction in the PRC2-deficient state (**Fig. 3D**). Leading-edge genes included canonical stress-response mediators such as *ATF4*, *DDIT3*, and C/EBP-family genes (**Supp. Fig. 3**).

We next asked whether this heightened stress response was accompanied by altered transcriptional programs supporting protein synthesis and RNA processing. A composite Translation/RNA activity index was generated by averaging standardized ssGSEA scores across pathways related to translation, ribosome biogenesis, and RNA processing; higher values therefore indicate greater relative activity of these programs within the dataset. HSV infection increased this index in PRC2-proficient cells (+0.54 units, *p*<0.001) but not PRC2-deficient cells (+0.09 units, *p*=0.27), producing a negative PRC2×infection interaction (−0.45, *p*<0.001; **Fig. 3E**). Thus, HSV infection elicited a coordinated increase in translation/RNA-associated transcriptional programs in PRC2-proficient cells that was blunted in the PRC2-deficient state. Together with the heightened ISR, this pattern is consistent with greater translational restriction in PRC2-deficient cells. Consistent with these findings, HSV G47Δ-treated S462-TY xenografts progressed and demonstrated limited viral distribution, with only patchy β-galactosidase staining within viable tumor 3 weeks after G47Δ treatment (**Fig. 3F**).

### BIRC5-regulated ICP34.5 restoration enhances antitumor activity of G47Δ

To test if restoring viral ISR antagonism increases HSV efficacy in H3K27me3-deficient MPNST, we engineered *BIRC5*-driven *ICP34.5* expression into the deleted *ICP34.5* loci of G47Δ (**Fig. 4A**; **Supp. Fig. 4**). *BIRC5*-driven reporter activity exceeded minimal-promoter activity, and the engineered vector attenuated HSV-induced eIF2α phosphorylation in S462-TY cells (**Supp. Fig. 4**).

**Figure 4:**
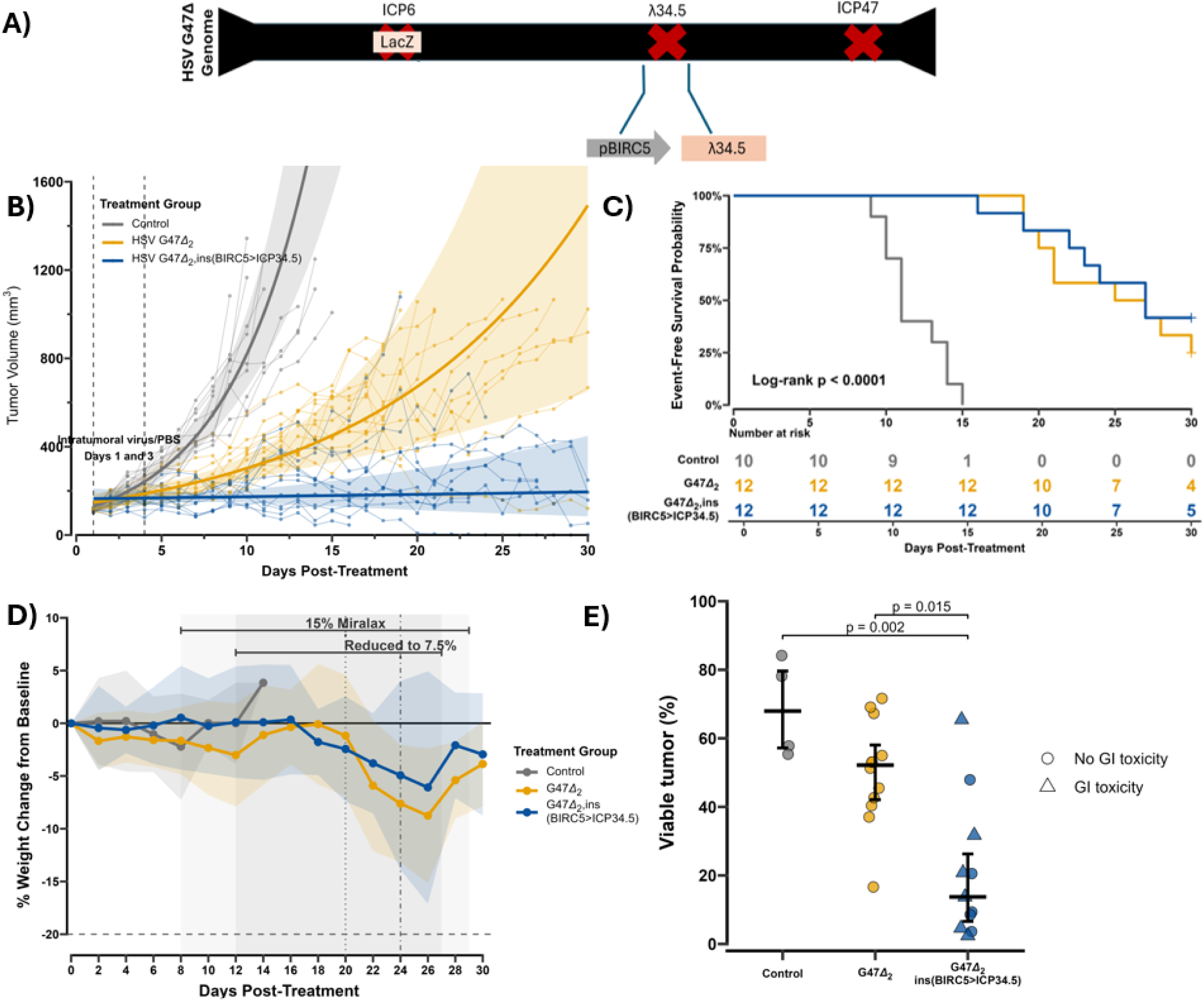
BIRC5-regulated restoration of ICP34.5 to counter the integrated stress response markedly enhances tumor control in H3K27me3-deficient MPNST. **A)** Schematic representation of an oncolytic HSV engineered to express ICP34.5 under the *BIRC5* promoter. The reverse-oriented SV40pA:pBIRC5>ICP34.5:rBGpA cassette was inserted into the two deleted ICP34.5 loci of HSV G47Δ via bacterial artificial chromosome (BAC) recombineering. The viruses utilized in vivo are annotated as HSV G47Δ_2_ and HSV G47Δ_2_,ins(BIRC5>ICP34.5). B) Tumor growth kinetics following intratumoral injection (Days 1 and 3) of PBS, HSV G47Δ_2_ (1.0×10^7^ PFU / dose), or HSV G47Δ_2_,ins(BIRC5>ICP34.5) (1.0×10^7^ PFU / dose). Tumor-growth rates were estimated using a linear mixed-effects model adjusted for sex. Control tumors exhibited an approximate daily tumor-volume change of 22.5% (95% CI, 17.4–27.8%), compared with 8.3% (95% CI, 4.5–12.2%) following HSV G47Δ_2_ and 0.6% (95% CI, −2.9–4.2%) following HSV G47Δ_2_,ins(BIRC5>ICP34.5). **C)** All-cause event-free survival. Both viral treatments significantly prolonged all-cause EFS relative to control (overall Log-rank *p*<0.001). During the trial, six G47Δ_2_,ins(BIRC5>ICP34.5) treated mice experienced unexpected gastrointestinal toxicity necessitating humane euthanasia. Pairwise log-rank testing revealed significant survival differences between control and both viral vectors (*p*<0.001), but no significant difference in all-cause survival between the two viral groups (*p*=0.58). **D)** Longitudinal weight monitoring following treatment. Weight is expressed as percent change from each animal’s pretreatment baseline; lines indicate group means and shaded ribbons indicate ±1 SD. The dashed horizontal line at −20% denotes the prespecified body-weight-loss euthanasia criterion. Following recognition of gastrointestinal toxicity, polyethylene glycol (Miralax) was introduced in the drinking water at 15% w/v and reduced to 7.5% four days later after dehydration-associated weight loss was observed. Because animals were enrolled on different calendar dates, horizontal brackets indicate the range of relative study day intervention and vertical lines indicate the median relative day. **E)** Viable tumor fraction quantified using a fixed QuPath pixel-classification model (Control, n=4; HSV G47Δ_2_, n=12; HSV G47Δ_2_,ins(BIRC5>ICP34.5), n=11). Viable tumor was reduced in the BIRC5>ICP34.5 group versus Control (Dunn-Holm *p*=0.002) and G47Δ_2_ (*p*=0.015). Triangles denote tumor samples extracted upon emergence of GI toxicity instead of tumor end-point or end-of-study.

In S462-TY xenografts, G47Δ_2_,ins(BIRC5>ICP34.5) markedly reduced tumor-growth kinetics relative to HSV G47Δ_2_ (pairwise slope comparison; Tukey-adjusted *p*=0.014). Model-estimated daily tumor-volume change was 22.5% (95% CI, 17.4–27.8%) in controls, 8.3% (4.5–12.2%) following HSV G47Δ_2_, and 0.6% (−2.9–4.2%) following HSV G47Δ_2_,ins(BIRC5>ICP34.5) (**Fig. 4B**). Exploratory extended-follow-up modeling of the ICP34.5-restored cohort identified a population tumor-trajectory change point at day 18.8 (approximate 95% profile-likelihood support interval, 17.0–20.2 days; **Supp. Fig. 5**). Only one mouse in the ICP34.5-restored cohort reached the prespecified tumor-volume endpoint at day +30, compared with 9/12 mice receiving parental HSV G47Δ_2_. In competing-risk analysis, the cumulative incidence of tumor endpoint differed significantly among virally treated groups (Gray’s test, *p*<0.001; **Supp. Fig. 5**).

### Unexpected gastrointestinal toxicity limited all-cause event-free survival of the ICP34.5-restored vector

Despite improved tumor control, all-cause EFS did not differ between the two viral groups on pairwise log-rank testing (*p*=0.58; **Fig. 4C**), reflecting six unexpected toxicity-related euthanasia events in the HSV G47Δ_2_,ins(BIRC5>ICP34.5) cohort. Clinical signs developed rapidly over 24 hours, and toxicity was characterized by gastric distention, ileus, and toxic megacolon, with median toxicity-related removal on day 23.5 (IQR, 22.2–26.2). Five of six toxicity-related removals had met the response criterion before removal. The median interval from confirmed response to toxicity-related removal was approximately 5 days. Following recognition of the toxicity, remaining animals received polyethylene glycol in drinking water (**Fig. 4D**), after which no additional toxicity-related removals occurred.^40^ The sole responder in the G47Δ_2_ group had colonic distention observed on necropsy at the time of end-of-study censoring, but demonstrated no clinical symptoms on laxative treatment. Tumors (N = 4 control, N = 12 G47Δ_2_, and N = 11 G47Δ_2_,ins(BIRC5>ICP34.5) as one subject had complete resolution of tumor burden) and matched livers (N = 4, 12, and 12, respectively) were assessed by immunohistochemistry. Consistent with tumor growth curves, the percent of viable tumor was reduced in the BIRC5>ICP34.5 group versus Control (Dunn-Holm *p*=0.002) and G47Δ_2_ (*p*=0.015), **Fig. 4E**. Ki67 staining correlated with viable tumor regions, while β-galactosidase staining negatively correlated with viable tumor regions (**Supp. Fig. 6**). Aside from the gastrointestinal system, no other gross end-organ damage was visualized on necropsy. On IHC, no liver samples demonstrated cytopathic effect nor β-galactosidase staining (**Table 2, Supp. Fig. 6**). No mice demonstrated clinical neurologic toxicity.

**Table 2:** Liver histopathologic and β-galactosidase assessment following intratumoral HSV treatment. Liver sections were evaluated for histologic cytopathic effect and β-galactosidase staining. No positive findings were observed in any treatment group.

**Liver histopathologic and β-galactosidase assessment**
| Liver endpoint | Control | G47$\Delta_2$ | G47$\Delta_2$,ins(BIRC5>ICP34.5) |
| --- | --- | --- | --- |
| Cytopathic effect | 0/4 | 0/12 | 0/12 |
| β-galactosidase staining | 0/4 | 0/12 | 0/12 |

Values indicate number positive / number assessed.

## DISCUSSION

H3K27me3-deficient MPNST lacks durable and effective treatment options, particularly for unresectable or metastatic disease. Surgery remains the curative cornerstone; radiotherapy benefits are uncertain, and chemotherapy is largely palliative.^10,42^ Our study illustrates how epigenetic vulnerabilities in H3K27me3-deficient MPNST can guide rational oncolytic HSV engineering for improved therapeutic efficacy.

HSV offers several features making it an attractive MPNST virotherapy platform, including extensive clinical experience with engineered vectors and neural crest tropism.^4,6,23^ This neural crest tropism may be particularly relevant to Schwann-cell-lineage malignancies. Our prior conditionally replicative adenovirus work required up to 50 MOI for complete *in-vitro* lysis, whereas HSV G47Δ produced cytotoxicity across human MPNST models at <1.0 MOI (**Fig. 1A**).^43^ More broadly, loss of repressive chromatin may favor infection by nuclear DNA viruses whose genomes are subject to host chromatin regulation (**Fig 2**). For HSV, the combination of this epigenetic vulnerability and intrinsic nervous system tropism provides a compelling rationale for therapeutic development in MPNST.

Prior studies have proposed that oncogenic MAPK signaling can attenuate PKR–eIF2α antiviral signaling, thereby contributing to tumor-selective replication of attenuated oncolytic viruses.^44^ Despite constitutive MAPK activation characteristic of MPNST, HSV infection induced eIF2α phosphorylation in our models (**Fig. 3B**), demonstrating that this antiviral effector axis remains intact. This observation provided the mechanistic basis for interrogating whether PRC2 loss alters the magnitude or transcriptional consequences of the antiviral stress response.

Heightened IFN-α, ISR, and apoptotic transcriptional programs in H3K27me3-deficient MPNST constitute a compensatory antiviral state, offsetting the transcriptional permissiveness conferred by PRC2 loss. This state may reflect direct transcriptional dysregulation or increased sensing of derepressed endogenous retroviral elements, as reported in H3K27-altered glioma.^45^ Notably, enrichment of apoptosis-associated pathways does not correspond to loss of viability, suggesting transcriptional priming without execution of cell death. Recurrent loss of *TP53* and *CDKN2A* in MPNST may contribute to this uncoupling.^42,46^ We propose exaggerated ISR signaling restricts viral protein synthesis and late assembly, separating efficient initial infection from sustained propagation. This may help explain the disparity between striking *in vitro* sensitivity to G47Δ and limited *in vivo* activity with HSV G47Δ, where successive rounds of infection must overcome physiologic barriers such as extracellular matrix and viral clearance, whereas cells are susceptible to repeated re-infection *in vitro*. Here, tumor-selective ICP34.5 restoration is not simply a strategy to increase HSV potency, but a rational means to overcome a host restriction mechanism that becomes consequential in H3K27me3-deficient tumors.

Conditional *ICP34.5* expression markedly enhanced tumor control in the aggressive S462-TY xenograft model, but the associated gastrointestinal toxicity requires careful consideration. The observed ileus and colonic distention phenotypically resembled the syndrome described by Khoury-Hanold et al., in which HSV-1 spread through sensory ganglia to the enteric nervous system in C57BL/6J mice, producing neutrophil-mediated loss of enteric neurons, impaired peristalsis, and lethal megacolon that could be prevented by laxative treatment.^40^ The predominant life-threatening manifestation of human visceral HSV dissemination is HSV hepatitis, which can rapidly progress to acute liver failure, coagulopathy, and death. In a series of 137 reported HSV-hepatitis cases, 84% presented with coagulopathy and 74% progressed to death or liver transplantation.^47^ In the present study, however, no histologic cytopathic effect or detectable β-galactosidase staining was identified in any liver from either viral treatment group, providing no histopathologic evidence of overt hepatic viral involvement (**Table 2**). The absence of hepatic injury does not necessarily exclude clinically relevant extrahepatic toxicity. Indeed, the severity and rapid progression of HSV hepatitis in reported human cases may obscure later or less immediately lethal manifestations of viral dissemination. An attenuated oncolytic HSV could therefore reveal toxicities not prominent in fulminant natural infection, including enteric dysmotility. Although uncommon in humans, alphaherpesvirus-associated paralytic ileus has been reported and resolved with conservative management.^48^ Colonic distention was also observed at necropsy in the sole parental G47Δ_2_ responder, despite the absence of clinical toxicity with laxative treatment. This suggests an HSV-dependent enteric mechanism can occur independently of restored ICP34.5 but rather be amplified by greater systemic viral exposure during highly productive replication. Prophylactic PEG may therefore have utility in future murine safety studies, while gastrointestinal dysmotility warrants prospective monitoring during extracranial clinical application of replication-competent oncolytic HSV vectors.

Transcriptional restriction of viral replication using BIRC5/survivin has precedent in conditionally replicating adenoviruses, in which survivin-regulated E1A expression produced preferential replication in malignant cells with limited replication in normal tissues, supporting the general feasibility of this targeting strategy.^26,30^ Prior MPNST work by Maldonado, et al evaluated *ICP34.5* restoration using the human midkine promoter, but it was evaluated in the PRC2-proficient STS26T, with just modest reduction in tumor growth rates, so safety in a productive infection context could not be evaluated.^49^ There are few other applications of *ICP34.5* expressing oncolytic HSV outside of the central nervous system. Given the lack of data, safer ISR-antagonizing strategies may be preferable for further development. Replacement of ICP34.5 with its *H. sapiens* cellular homolog PPP1R15A/GADD34 preserves eIF2α dephosphorylation while eliminating ICP34.5 neurovirulence functions such as Beclin-1 binding. A GADD34-expressing oncolytic HSV retained antitumor efficacy comparable with an ICP34.5-restored vector while demonstrating improved tolerability in susceptible mice.^20,24,25^ Such approaches may preserve the replicative advantage identified here while reducing off-target toxicity for further development in MPNST.

Our study has important limitations that can inform additional future investigations. This study was designed primarily to query mechanisms governing MPNST susceptibility to oncolytic HSV. For future toxicology studies, orthotopic peripheral-nerve models will be important to evaluate local neurotoxicity not captured by flank xenografts. Moreover, our *in vivo* studies did not directly test whether ICP34.5-restored oncolytic HSV has differential activity across PRC2 states, which is best assessed in an isogenic model given intrinsic differences in growth rates and innate anti-viral responses between distinct models. We attempted different strategies, but each had limitations. Systemic doxycycline produced only patchy H3K27me3 restoration in TY5005 xenografts, as assessed by IHC, while C57BL/6-derived syngeneic models were poorly permissive to HSV, as assessed by flow cytometry, limiting both approaches. BALB/c mice are comparatively susceptible to HSV-1 and have been used for preclinical evaluation of *ICP34.5*-regulated oncolytic HSV.^39^ Thus BALB/c-compatible syngeneic MPNST models, which have been developed,^50^ may enable isogenic biomarker testing and adaptive immune interactions in an immunocompetent host.

The implications of this work may extend beyond MPNST. Global H3K27me3 loss is also a defining feature of diffuse midline glioma and posterior fossa group A ependymoma, providing a rationale to test whether this epigenetic state similarly predicts oncolytic HSV susceptibility in these diseases. The clinical experience with CAN-3110/rQNestin34.5v.2 demonstrates that tumor-regulated *ICP34.5* expression can be administered safely to patients with high-grade glioma, although therapeutic benefit remains heterogeneous.^22,23^ Biomarker selection based on tumor epigenetic state may therefore provide one strategy for refining deployment of such vectors. Conversely, pharmacologic PRC2 inhibition could potentially sensitize PRC2-proficient tumors to oncolytic HSV, although effects on normal-tissue permissiveness would require careful evaluation. These findings also suggest broader translational opportunities within the Schwann-cell lineage. In MPNST, oncolytic HSV could ultimately be integrated with maintenance strategies targeting the same H3K27me3-deficient biology, including strategies under active evaluation (e.g. NCT06693284). More attenuated or replication-restricted HSV platforms may also warrant investigation in nonmalignant Schwann-cell tumors, including plexiform neurofibromas and schwannomas, where treatment goals center on morbidity reduction rather than tumor eradication.^51,52^ Although beyond the scope of the present study, these directions illustrate how lineage susceptibility to HSV could be leveraged across a broader spectrum of peripheral nerve sheath disease.

## CONCLUSIONS

Our work identifies H3K27me3 loss in high-grade MPNST as a therapeutically exploitable vulnerability to oncolytic HSV, while revealing a compensatory integrated stress response that limits productive viral propagation. Engineering HSV to overcome this second antiviral barrier markedly enhanced antitumor activity. These findings provide a framework for biomarker-driven, gene-enhanced virotherapy in MPNST and demonstrate how tumor-specific defects in antiviral restriction can guide rational viral engineering.

## REQUIRED STATEMENTS

### Ethics

Animal experiments were conducted in accordance with federal and institutional guidelines and approved by the Institutional Animal Care and Use Committee (IACUC) at the University of Minnesota (Protocol ID: 2407-42182A). The university maintains an AAALAC-accredited animal facility and operates under NIH Animal Welfare Assurance number D16-00288 (A3456-01) and USDA Registration Number 41-R-0005. Mice were maintained under BSL2 containment.

### Funding

This work was supported by funding from the 2025 Children’s Tumor Foundation Drug Discovery Initiative, a gift from Humor to Fight the Tumor, a gift from the Rick Bock Foundation, and the University of Minnesota Department of Pediatrics.

### Conflict of Interest

Dr. Largaespada is the co-founder and co-owner of NeoClone Biotechnologies, Inc., Discovery Genomics, Inc. (recently acquired by Immunsoft, Inc.), B-MoGen Biotechnologies, Inc. (recently acquired by the Biotechne corporation), and Luminary Therapeutics, Inc. He consults for Styx Biotechnologies and Genentech, Inc., which has funded some of his research. DayOne Biopharmaceuticals is also funding some of his lab’s research. The business of all these companies is unrelated to the contents of this manuscript. All other authors have no conflicts to disclose.

### Authorship

- Conceptualization: RTG
- Data curation: RTG, MS, LL, BGS, AH, SP, EW, DM, CB, LF, ZS, WH
- Formal analysis: RTG, MS, LL, BGS, AH, SP, EW, DM, KW, DAL
- Funding acquisition: RTG, DAL
- Investigation: RTG, MS, LL, BGS, AH, SP, EW, CB, DM, CS, MS, LF, CJL, KW, AL, ZS, WH, DAL
- Methodology: RTG, MS, LL, BGS, LF, DM, CB, CS, MS, CJD, KW, AL, ZS, WH, DAL
- Project administration: RTG
- Resources: RTG, LL, BGS, CS, MS, LF, CJD, KW, AL, ZS, WH, DAL
- Supervision: RTG
- Validation: RTG, LL, BGS, DM, KW
- Visualization: RTG, LL, BGS, AH, DM
- Writing – original draft: RTG, BGS, DM
- Writing – review & editing: RTG, MS, LL, BGS, SP, EW, DM, CS, MS, CJD, KW, AL, ZS, WH, DAL

### Data Availability

Raw CUT&RUN sequencing data and processed coverage tracks have been deposited in the Gene Expression Omnibus (GEO) under accession number GSE329989. Raw RNA-sequencing data and processed count matrices have been deposited in the Gene Expression Omnibus (GEO) under accession number GSE329232. Raw immunohistochemistry files are hosted on Histowiz and available at: https://app.histowiz.com/shared_groups/2683a825-544c-42cf-af50-fe7a59c17450/slides **[IHC]** https://app.histowiz.com/shared_groups/8bab4411-6f50-4b53-81a3-fd0f9cb02a38/slides **[H&E].**

## ACKNOWLEDGEMENTS

We kindly acknowledge Dr. Jianfang Ning who gifted an aliquot of HSV G47Δ for this work. We kindly acknowledge cell lines gifts from Dr. Christine Pratilas (MPNST724, JW18, JH-2-002) and Dr. Nancy Ratner (S462-TY, STS26T, ST88-14). Dr. Galvin sincerely acknowledges Dr. Christopher Moertel and Dr. Brenda Weigel for their mentorship. We kindly acknowledge services provided by the University of Minnesota Genomics Center, Minnesota Supercomputing Institute, the University of Minnesota Flow Cytometry Resource, and the University of Minnesota Research Animal Resources. This work was supported by funding from Humor to Fight the Tumor, Children’s Tumor Foundation Drug Discovery Initiative, the Rick Bock Foundation, and the University of Minnesota Department of Pediatrics.

## Notes

https://www.ncbi.nlm.nih.gov/geo/query/acc.cgi?acc=GSE329232

https://www.ncbi.nlm.nih.gov/geo/query/acc.cgi?acc=GSE329989

