## Supplementary Figures 1-6 for "Biomarker-driven and gene-enhanced viral therapy for malignant peripheral nerve sheath tumor"

### SUPPLEMENTARY FIGURE 1

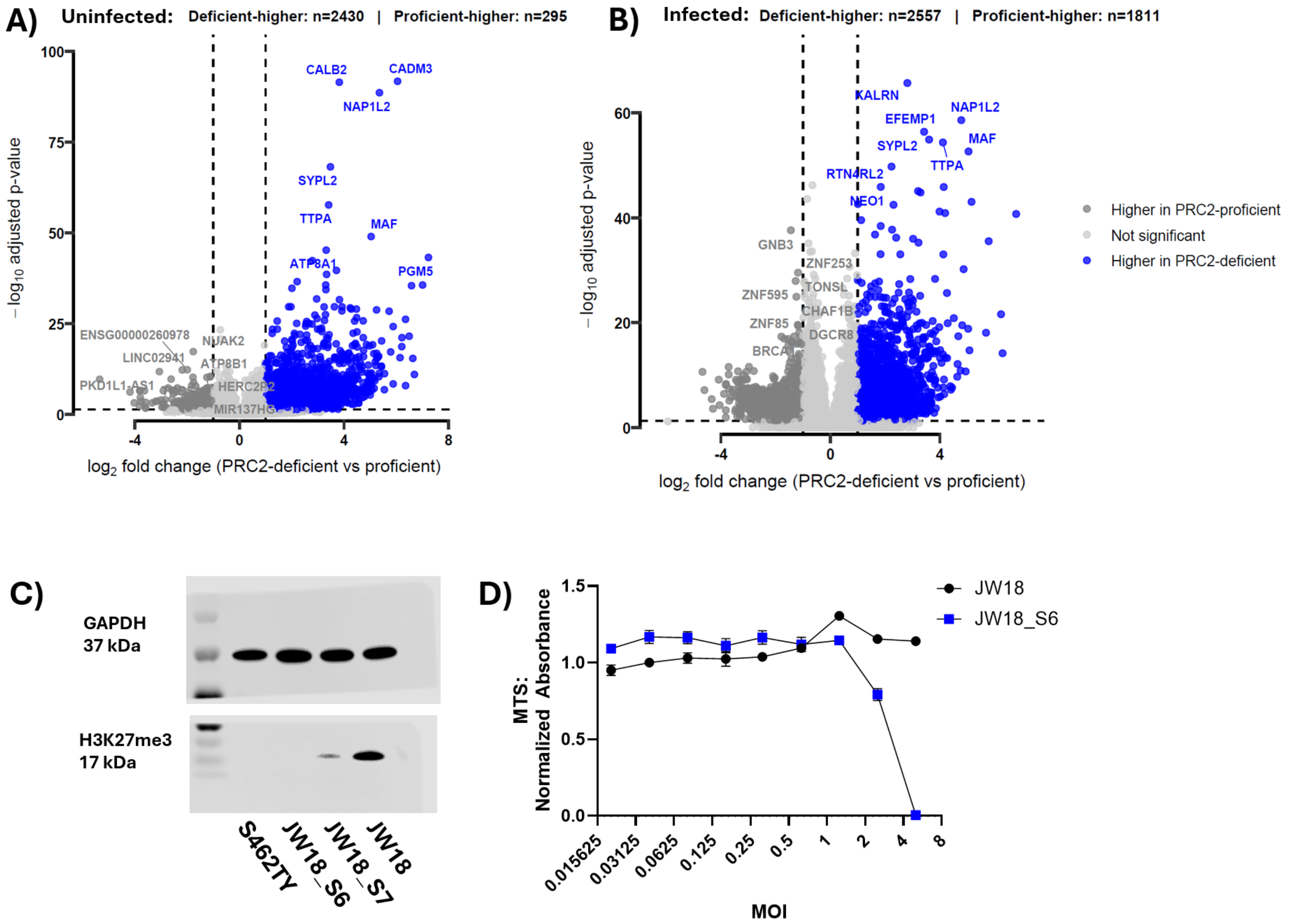


**Supplementary Figure 1: Global transcriptional response by PRC2 phenotype and validation of H3K27me3-dependent viral sensitivity in a murine MPNST model.** A-B) Volcano plots detailing global differential gene expression from RNA-sequencing of PRC2 modular systems in uninfected (A) and HSV-infected (B) conditions. Positive log2 fold changes (x-axis) represent genes significantly upregulated in the PRC2-deficient phenotype (blue), while negative values represent genes upregulated in the PRC2-proficient phenotype (grey). Notably, while the number of overexpressed genes in the PRC2-deficient context remains relatively stable regardless of infection status (n=2430 uninfected vs. n=2560 infected), the PRC2-proficient context reveals a marked increase in the fraction of differentially expressed genes in response to the virus (n=295 uninfected vs. n=1815 infected). C) Western blot validation of CRISPR/Cas9-mediated *Suz12* knockout in the murine NPcis MPNST cell line JW18 ($Tp53^{-/-}$ and $Nf1^{-/-}$). The resulting derivative clone, JW18_S6, demonstrates suppression of global H3K27me3 compared to the PRC2-proficient parental line. GAPDH serves as a loading control and lysate from S462-TY as a negative control. D) In vitro cytotoxicity of HSV G47Δ against the murine isogenic pair, assessed via MTS assay. Consistent with human MPNST models and primary murine Schwann cell models (Figure 1), the H3K27me3-deficient murine clone (JW18_S6) exhibits enhanced sensitivity to virus-mediated cell death compared to the parental JW18 line. Murine syngeneic cell lines demonstrate intrinsically low permissiveness to HSV replication, necessitating substantially higher viral doses (MOI > 1) to achieve lytic efficacy compared to human MPNST models.

**SUPPLEMENTARY FIGURE 2**

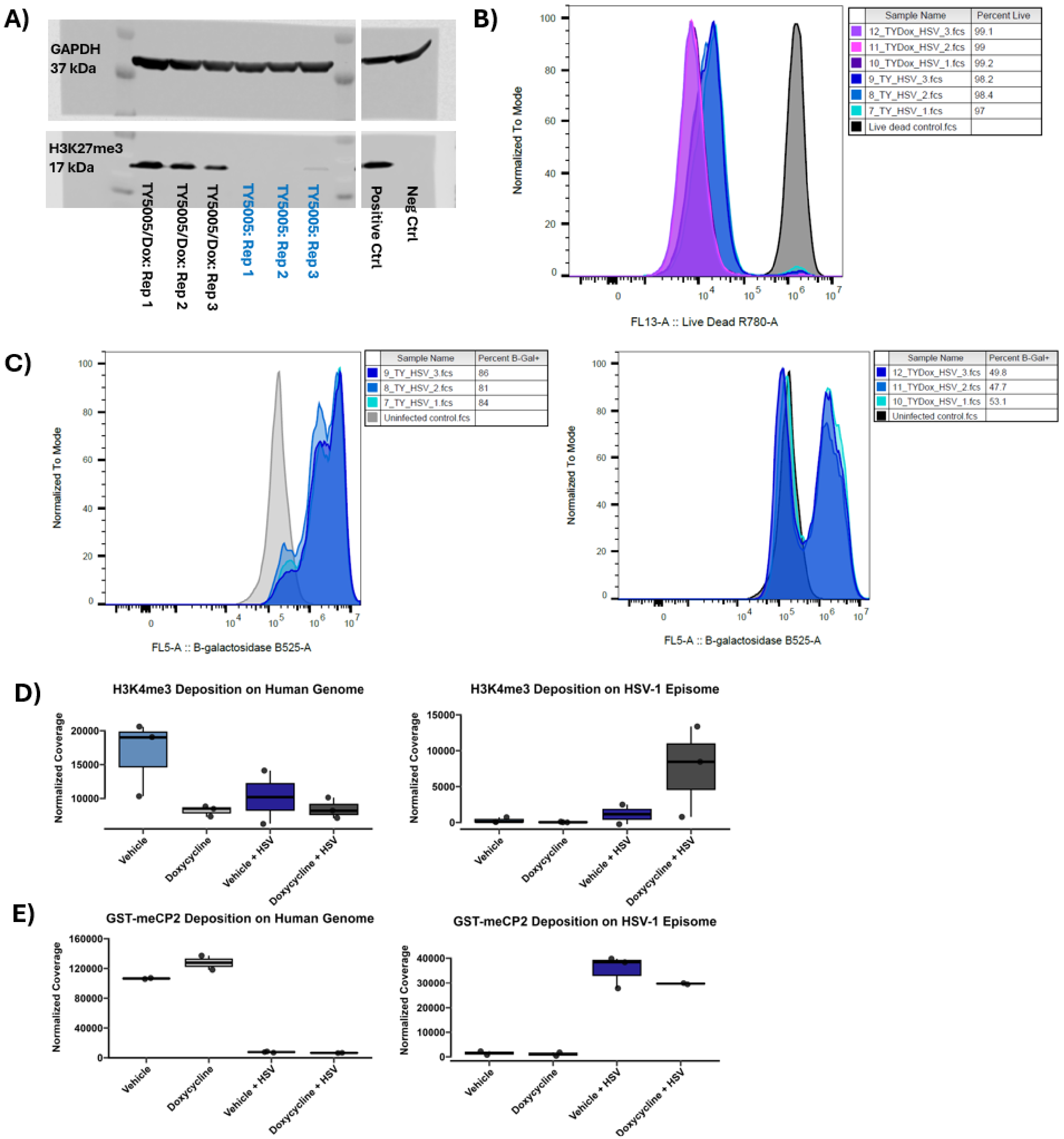


**Supplementary Figure 2: CUT&RUN sample validation and exploratory epigenetic profiling of H3K4me3 and GST-meCP2.**

A-C) Quality control metrics for the TY5005 biological replicates submitted for CUT&RUN analysis (see Figure 2). A) Western blot analysis confirming the expected H3K27me3 status across all individual biological replicates prior to processing. Consistent H3K27me3 restoration is observed in the doxycycline-induced (TY5005/Dox) replicates compared to the uninduced controls. β-actin serves as a loading control. B) Flow cytometry viability assessment (eBioscience™ Fixable Viability Dye eFluor™ 780) demonstrates that all biological replicates retained high cell viability (>97% live cells) at the time of sample acquisition. C) Flow cytometry quantification of viral β-galactosidase expression (Spider-βGal, Dojindo) at the time of sample collection. Highlighting the differential kinetics of viral spread between the phenotypes, H3K27me3-deficient replicates achieved >80% infectivity, whereas H3K27me3-proficient replicates achieved approximately 50% infectivity at the collection time point. D-E) Exploratory CUT&RUN quantification of H3K4me3 (D) and GST-meCP2 (E) deposition on the human genome (left) and HSV-1 episome (right). Y-axes represent total spike-in scaled occupancy mathematically background-subtracted against matched IgG negative controls. Following infection, H3K4me3 and GST-meCP2 signals decreased across the host genome while increasing over the HSV-1 genome. Assay-specific technical QC failures were excluded from quantitative analysis: Vehicle+HSV replicate 3 for H3K27me3 and H3K4me3 because of an aberrant shared IgG control; Vehicle replicate 3 GST-meCP2 because of apparent contamination (~$1\times10^6$ HSV-aligned reads in an uninfected sample); Doxycycline replicate 2 GST-meCP2 because of an extreme spike-in count (~$14\times10^6$ reads); and Doxycycline+HSV replicate 3 GST-meCP2 because of spike-in dropout. Unaffected assays from these biological replicates were retained.

**SUPPLEMENTARY FIGURE 3**

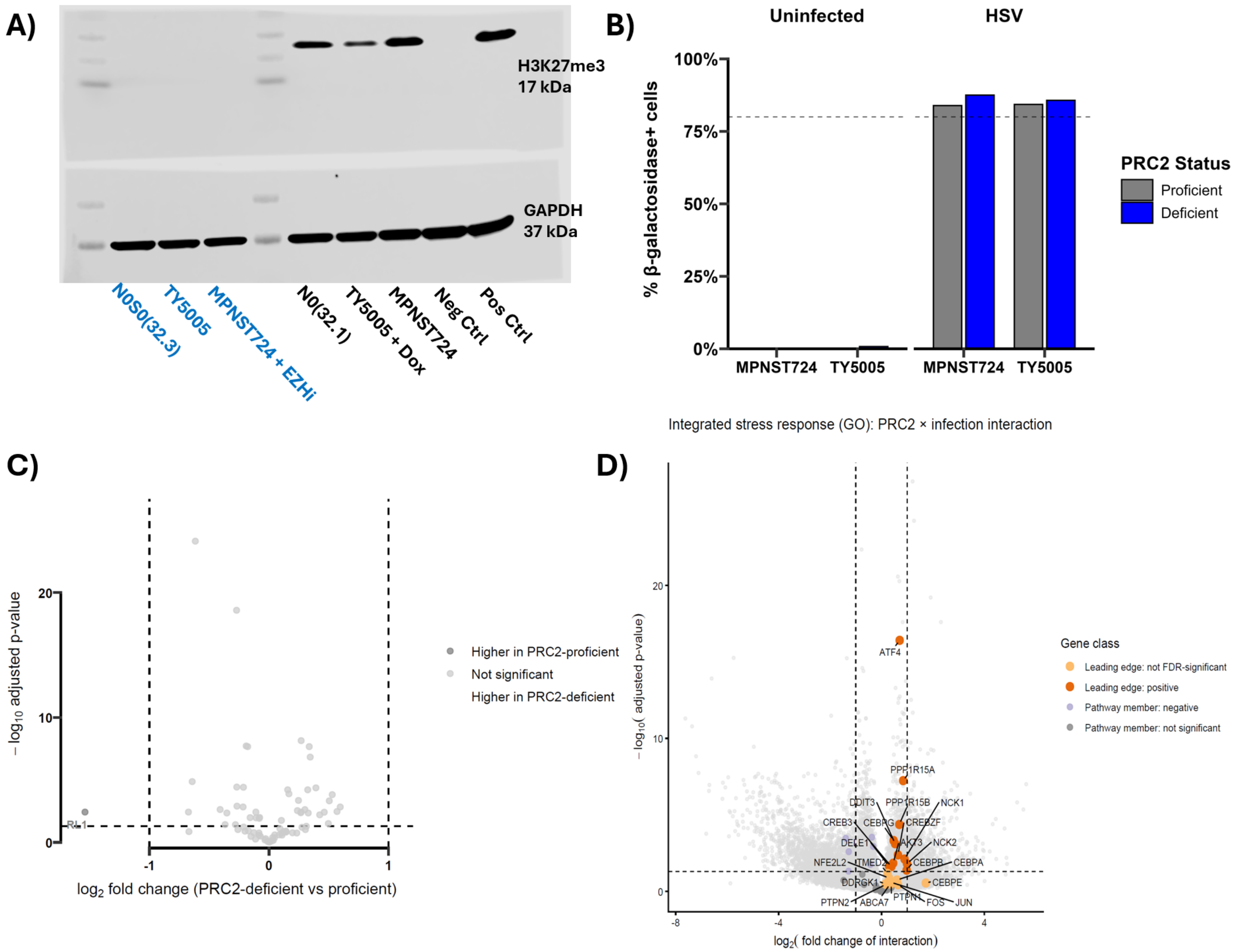


**Supplementary Figure 3: Quality control metrics and leading-edge gene analysis for the global transcriptomic profiling of MPNST models following HSV infection.** A-C) Quality control validation for the RNA-sequencing cohort presented in Figure 3. **A)** Representative immunoblot confirming the expected modulation of H3K27me3 across the three paired MPNST model systems used for RNA sequencing. Because the complete sequencing cohort comprised 72 independently processed RNA samples, immunoblot validation was performed on one representative biological replicate per condition. β-actin served as a loading control. **B)** Viral-burden quality control at the time of RNA-sequencing sample acquisition. Six independent biological replicates were maintained for each PRC2-status and infection condition. Because PRC2-proficient and PRC2-deficient cells exhibited different HSV replication kinetics, model-specific MOIs and harvest times were empirically optimized to achieve comparable viral spread without cytotoxicity. Beginning 24 h after infection, small aliquots from the six biological replicates within each condition were pooled solely for infection-quality-control assessment and monitored at 24-h intervals; biological replicates remained separate for subsequent RNA extraction and sequencing. Infected samples were harvested when the pooled condition reached a target viral burden of >80% infected cells (dashed line), with matched uninfected controls collected at the corresponding elapsed time. Collection occurred 36–72 hr after infection depending on model and PRC2 state. For the TY5005 and MPNST724 systems, infection was quantified using Spider-βGal detection of viral β-galactosidase expression. Because constitutive GFP expression in the N32 system interfered with fluorescence-based Spider-βGal detection, N32 infection was assessed semi-quantitatively by visual estimation of X-gal-positive cells. **C)** Differential expression of HSV transcripts between PRC2-deficient and PRC2-proficient infected samples. Most viral genes showed no significant PRC2-associated differential expression, indicating broadly similar relative viral transcriptional profiles across conditions. D) Volcano plot highlighting host genes belonging to the leading edge of the significant GO ISR pathway enrichment for the PRC2 × infection contrast (identified in Figure 3). Key ISR mediators (e.g., *ATF4*, *DDIT3, CEBP/C/EBP* family members) are prominently enriched.

#### SUPPLEMENTARY FIGURE 4

**A)**

Path of agarose gel image

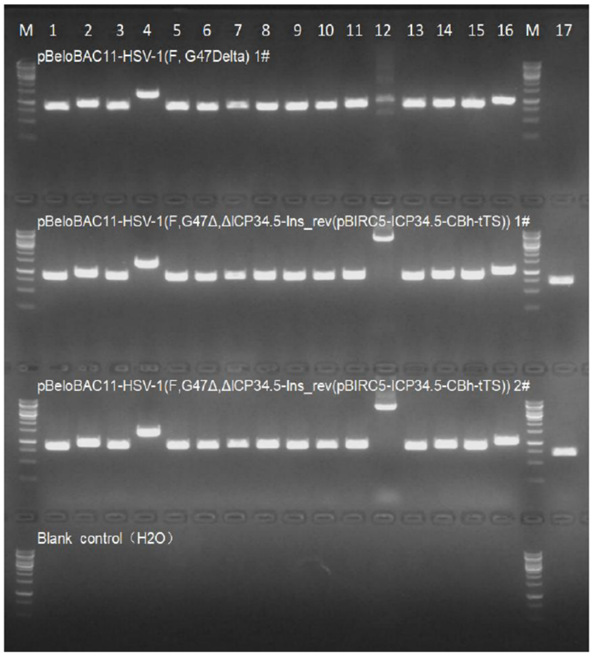


| Lane | Primer | | Length(bp |
|---|---|---|---|
| 1 | F1 | CCAACGAAAAGCGCGGGAACGG | 853 |
| | R1 | CCTGACGAGCGACAGGCGGCCG | 853 |
| 2 | F2 | GGCGCTGGTCTTGCGGCCGATC | 951 |
| | R2 | AAATTCAGACGGCAAACGACTG | 951 |
| 3 | F3 | GCACGCGTACAGCGCCCGCCAC | 880 |
| | R3 | CGAACTGTTTGGCGTGCAGATG | 880 |
| 4 | F4 | CAGTTCACCCGTGCACCGCTGG | 1392 |
| | R4 | GCAGCTGGAGGTGTAGTAGTTG | 1392 |
| 5 | F5 | GGCAGAGGTAGGCGTAGCTTTC | 883 |
| | R5 | CTGGTCGAGTGGTTGGATCGCG | 883 |
| 6 | F6 | GTAATGGCCTCGCGGTCGTAAC | 871 |
| | R6 | CAAAGATCGTGTTATGGATAAG | 871 |
| 7 | F7 | GTAATCATAACCACGCCCAGCG | 897 |
| | R7 | GCTATTTTACGCCTCGGCTGTC | 897 |
| 8 | F8 | CCGACATGGTGGCGATAGACGC | 895 |
| | R8 | TGCGGGAACACGCGTACAGCCG | 895 |
| 9 | F9 | GTTACGCCCGATCGAACGGAAC | 875 |
| | R9 | GCAGAGTCTGGTCCTCCCACAG | 875 |
| 10 | F10 | CAACCACGGTCACGCTTCGGTG | 867 |
| | R10 | GCGTGGTGGCGTCGATGGTGTC | 867 |
| 11 | F11 | GAAGATGGCCGCGTGGAACTGC | 895 |
| | R11 | CCGCGGAGCCATCGGCGTCTTC | 895 |
| 12 | F12 | AAGAATATCATTAGCATGCACG | 1011 |
| | R12 | GGTGTTGTTACACCTGTGGCAG | 1011 |
| 13 | F13 | CTATTTGGTGGGTGGTTGGTGC | 857 |
| | R13 | CCGATGACTCCCAAGAGGTGAC | 857 |
| 14 | F14 | GGTCCTCCGGGACGTTTTCTGG | 895 |
| | R14 | ACTAAGATCACACTCCGGGCCG | 895 |
| 15 | F15 | GACCTCCAGGTCCCTGGGAGAC | 894 |
| | R15 | CTGGAGGACGCCATCGTGCTGC | 894 |
| 16 | F16 | CGCGGGCCCGAGCGTACGCTCC | 1185 |
| | R16 | TTAGGGCGAAGTGCGAGCACTG | 1185 |
| 17 | F17 | AAGAATATCATTAGCATGCACG | 701 |
| | R17 | TCGTCCCTGCCCACACCTAGCG | 701 |

Marker

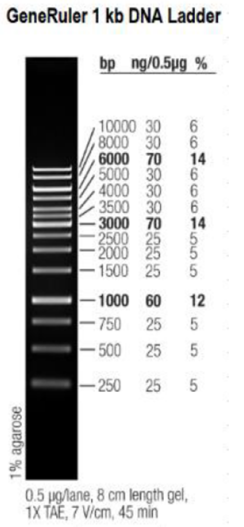


**B)**

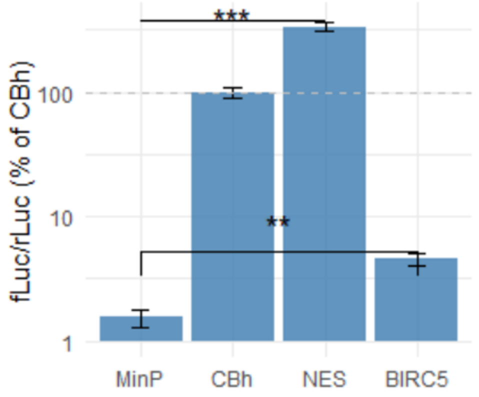


**C)**

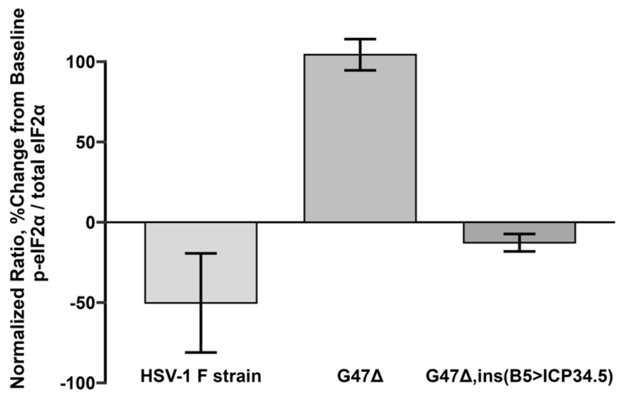


**Supplementary Figure 4: Validation and functional characterization of HSV G47$\Delta_2$,ins(BIRC5>ICP34.5).**

**A)** PCR validation of engineered HSV BACs. Lane 12 spans the modified *ICP34.5* locus; the parental HSV G47$\Delta_2$ BAC yields the expected 1,011-bp amplicon, whereas insertion of the reverse-oriented SV40pA:pBIRC5>ICP34.5:rBGpA-CBh>tTS:SV40pA cassette increases the expected amplicon size to 5,078 bp. Bands at the remaining assayed loci were consistent with their predicted sizes, supporting the expected BAC architecture. **B)** Promoter activity in S462-TY cells assessed by dual-luciferase reporter assay. Firefly luciferase activity driven by MinP, CBh, NES, or BIRC5 was normalized to cotransfected Renilla luciferase and expressed relative to the CBh>Luc2 positive control. BIRC5-driven activity was significantly greater than MinP

($p$<0.01, Welch's $t$-test) but lower than that observed with the NES regulatory element. **C)** Ratio of phosphorylated to total eIF2α in S462-TY cells infected with wild-type HSV-1 F, HSV G47$\Delta_2$, or HSV G47$\Delta_2$,ins(BIRC5>ICP34.5), quantified by immunoblotting in $n$ = 3 independent biological replicates and normalized to matched uninfected controls. Error bars represent SD. Increased eIF2α phosphorylation was observed with HSV G47$\Delta_2$ but was attenuated with HSV-1 F and HSV G47$\Delta_2$,ins(BIRC5>ICP34.5), consistent with functional ICP34.5-mediated antagonism of eIF2α phosphorylation in the engineered vector.

**SUPPLEMENTARY FIGURE 5**

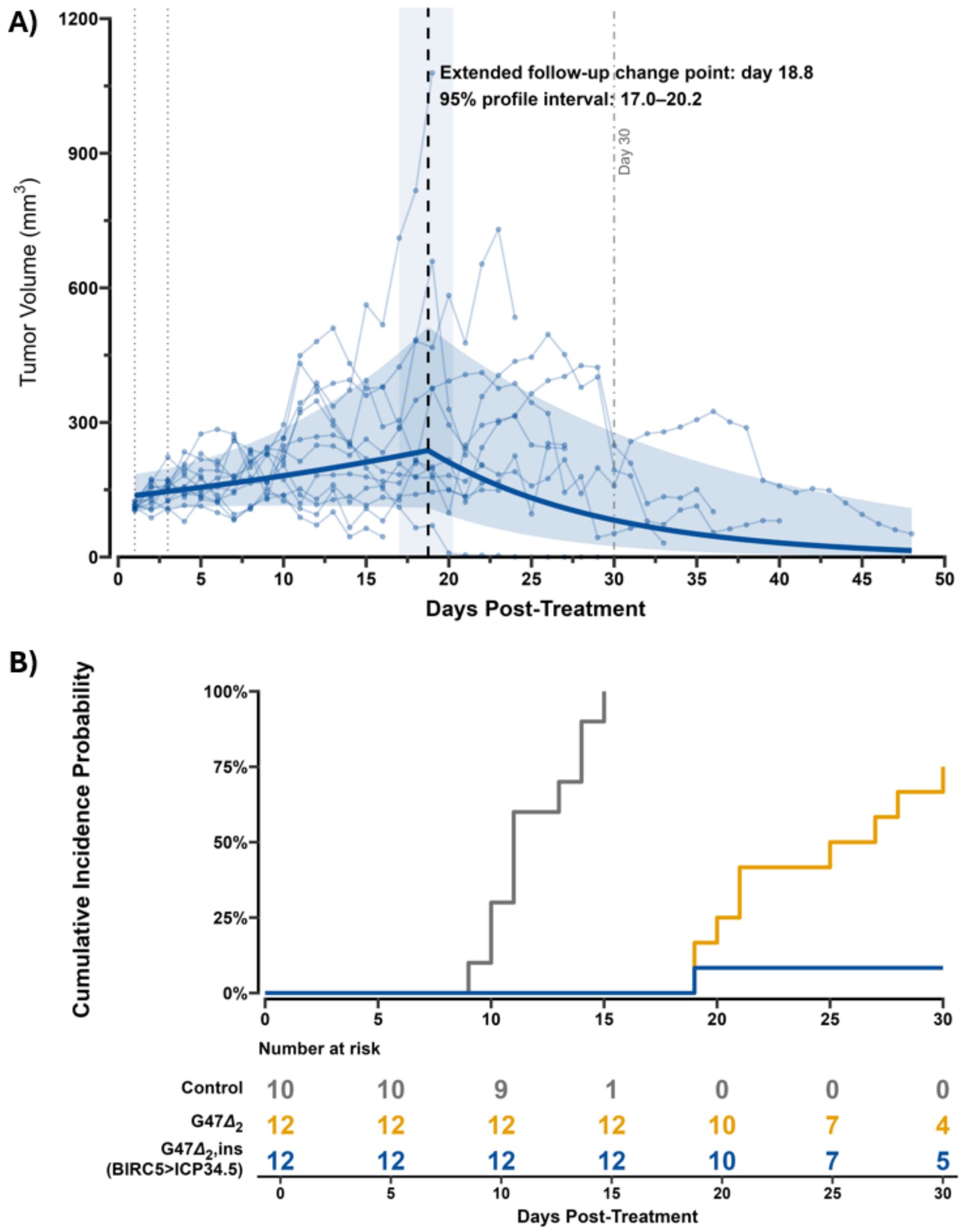


**Supplementary Figure 5: Extended response kinetics and tumor-endpoint incidence following HSV $G47\Delta_2$,ins(BIRC5>ICP34.5) therapy.**

**A)** Extended longitudinal tumor-volume trajectories for mice treated with HSV $G47\Delta_2$,ins(BIRC5>ICP34.5), including observations recorded beyond the prespecified day-30 analysis window. Thin lines and points represent individual animals; the solid line and shaded band represent the fitted population trajectory and 95% confidence interval from an exploratory piecewise linear mixed-effects model of log1p-transformed tumor volume. Candidate change points were evaluated by maximum-likelihood fitting, with the minimum-AIC model identifying a transition at day 18.8 after treatment (approximate 95% profile-likelihood interval, 17.0–20.2 days).

The dashed vertical line denotes the estimated change point, the shaded vertical region its profile interval, dotted lines indicate treatment administration on days 1 and 4, and the gray vertical line marks day 30. Because follow-up beyond day 30 was contributed only by prolonged survivors, the extended analysis was considered exploratory. **B)** Cumulative incidence of tumor endpoint in the presence of competing non-tumor events. The cumulative incidence of reaching the prespecified tumor-volume endpoint differed significantly across treatment groups (Gray's test, $p$<0.001), with only one mouse in the HSV G47$\Delta_2$,ins(BIRC5>ICP34.5) cohort reaching the tumor endpoint.

**SUPPLEMENTARY FIGURE 6**

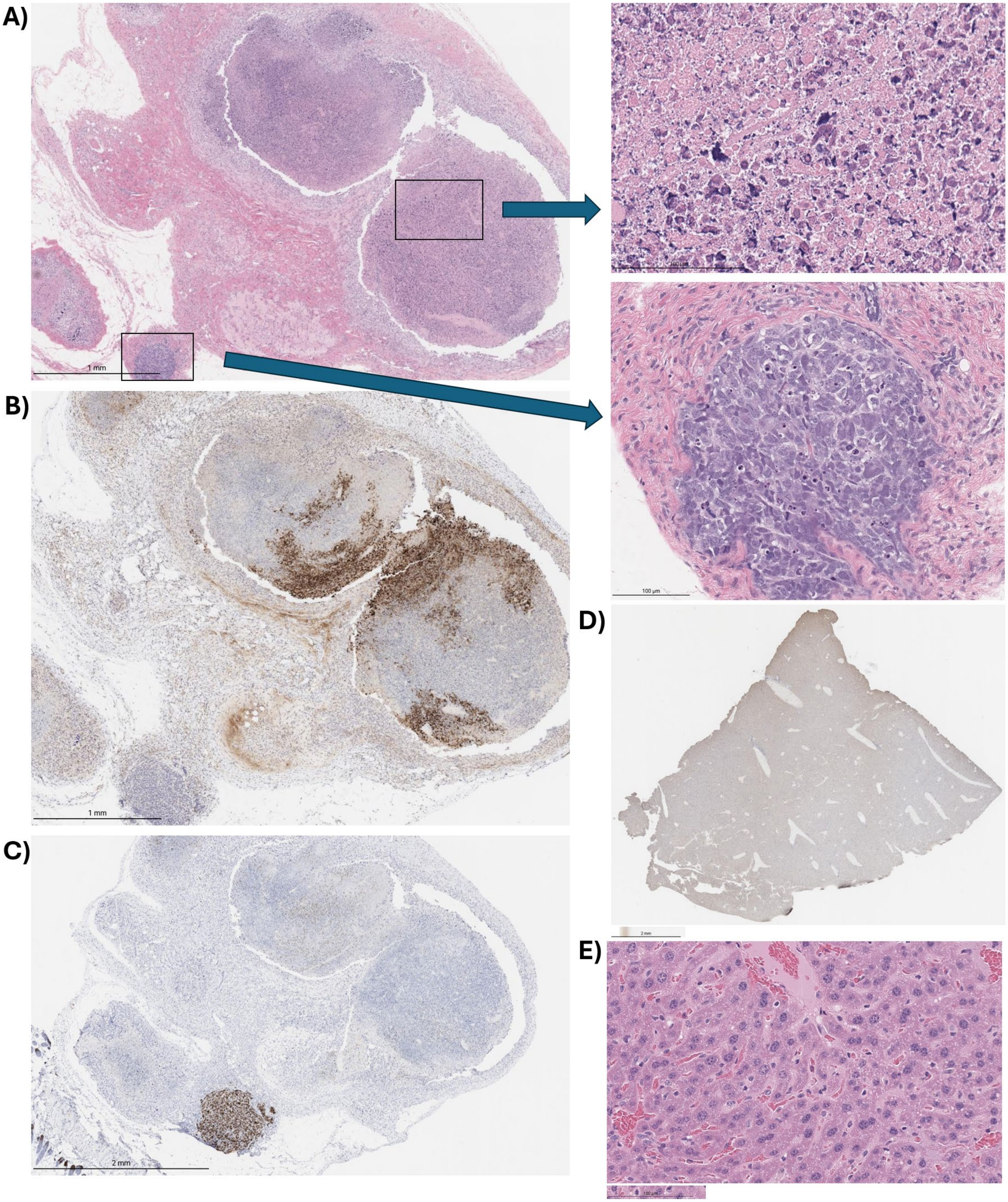


**Supplementary Figure 6. Representative histopathology and immunohistochemistry from HSV**

**G47$\Delta_2$,ins(BIRC5>ICP34.5)-treated S462-TY xenografts.** A–C) Matched tumor sections from mouse A2.4F, censored on day 32 after study enrollment (tumor volume, 52.7 $mm^3$). This tumor, together with a control tumor, was used to train the pixel-classification model for viable tumor, degenerative tumor, and necrotic/fibrotic tissue. A) H&E demonstrates heterogeneous treatment-associated morphology, including extensive necrosis and large nodules with degenerative/cytopathic features and marked morphologic atypia (upper inset; scale bar, 100 μm). A small focus of morphologically preserved S462-TY tumor is shown in the lower inset (scale bar, 100 μm). Whole-slide scale bar, 1 mm. B) β-galactosidase staining is most prominent within regions exhibiting degenerative/cytopathic morphology. Scale bar, 1 mm. C) Ki67 staining demonstrates focal proliferative activity within residual viable tumor. Scale bar, 2 mm. D–E) Representative liver histology from mouse A4.2F, euthanized on day 27 because of gastrointestinal toxicity. D) Whole-slide β-galactosidase staining demonstrates no detectable hepatic β-galactosidase signal. Scale bar, 2 mm. E) Representative H&E demonstrates preserved hepatic architecture without overt cytopathic change. Scale bar, 100 μm.
