## Supplementary Methods for "Biomarker-driven and gene-enhanced viral therapy for malignant peripheral nerve sheath tumor"

1. **Murine syngeneic model with loss of *Suz12***

The NPcis-derived syngeneic MPNST cell line JW18 (*Tp53*$^{-/-}$, *Nf1*$^{-/-}$) was edited using CRISPR/Cas9 ribonucleoprotein nucleofection. Three *Suz12*-targeting sgRNAs were used: 5′-CGCCUCAGAAGCACGGCGGU-3′, 5′-GGUCCGGGGGAGGCGGCUUC-3′, and 5′-GCUUCGGGCGGCAAAUCCGG-3′. RNPs were delivered using P3 Nucleofector solution and pulse code DS-150. Editing was screened by PCR using GC-rich Taq polymerase, forward primer 5′-GGCGAGCGGTTGGTGTAG-3′, and reverse primer 5′-CACCAACTCGTCTCGGGAT-3′, followed by Sanger sequencing with the reverse primer. Single-cell clones were isolated in 96-well plates.

2. **HSV-1(F) Vector Engineering**

Bacterial Strains and Culture Conditions All modifications were performed using the pBeloBAC11-HSV-1(F) Bacterial Artificial Chromosome (BAC) propagated in *E. coli*. The E. coli backbone in this construct interrupts the viral *UL23* gene (Thymidine Kinase). Cells were cultured in Luria-Bertani (LB) broth or on LB agar plates. Antibiotics were used for selection: Chloramphenicol (Chl) for BAC maintenance, Ampicillin (Amp), Kanamycin (Kan), and Tetracycline (Tet) for selection cassettes and helper plasmids. L-arabinose was used to induce recombinase expression.

Generation of pBeloBAC11-HSV-1(F, ΔICP47_363bp, ΔICP34.5, ICP6::LacZ)

*ICP47 deletion*: The 363 base pair coding region of ICP47 from the HSV-1 F strain (Accession GU734771.1) was deleted per the HSV G47Δ design.[6] Prior to modification, the pBeloBAC11-HSV-1(F) backbone was validated via PCR to confirm the integrity of the ICP47 flanking regions (homologous arms) using primers CGCGGGCCCGAGCGTACGCTCC (Forward) and TCCTGGACAACATGCGGGTTGG (Reverse).

First, a ccdB-amp selection cassette was introduced via λ Red recombination utilizing the temperature-sensitive helper plasmid pSC101-ccdA-gbaA.[31] The targeting fragment was generated via homology arms corresponding to the sequences flanking the ICP47 locus:
5' Homology Arm:
ACGAGGACGTTCCTCCTGCGGGAAGGCACGAACGCGGGTGAGCCCCCTCCTCCGCCCCCG
3' Homology Arm:
ACGGGGGGAACCACCCTCCGCCCAGAAACTTGGGCGATGGTCGTACCCGGGACTCAACGG

Recombinants were selected on LB agar with Chl, Amp, Tet, and L-arabinose at 30°C. Clones underwent PCR using wildtype primers and recombination-specific primers AGTTCGCCAGTTAATAGTTTGC (Forward); TTAGGGCGAAGTGCGAGCACTG (Reverse) and products were analyzed via gel electrophoresis. Successful clones were cryopreserved. The target fragment in the selected clone was subjected to confirmatory Sanger sequencing.

Next, the ccdB-amp cassette was removed through a second round of recombination. Following induction, cells were plated on LB agar containing only Chloramphenicol and incubated at 37°C. Clones were screened via gel electrophoresis using updated recombination-specific primers CGCGGGCCCGAGCGTACGCTCC (Forward); TTAGGGCGAAGTGCGAGCACTG (Reverse). Successful recombination was identified by clones with absent wildtype bands and present recombination-specific bands (**Fig. 1**). The target region underwent confirmatory Sanger sequencing.

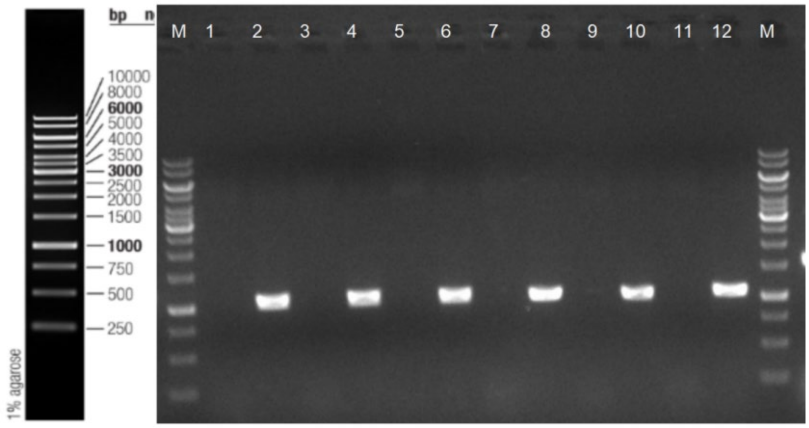


**Figure 1**: Colony PCR Gel Electrophoresis, *ICP47* deletion. Odd lanes represent wild-type specific sites, even lanes represent recombination-specific sites. All 6 clones demonstrate successful recombination.

*ICP34.5 deletion*:

The two copies of the ICP34.5 (RL1) region were targeted for markerless deletion via a two-step site-specific recombination process, per the HSV G47Δ design.[32] Target regions within the ICP34.5 loci were replaced with FRT-flanked kanamycin and a Rox-flanked ampicillin selection cassettes, respectively. The targeting fragments were generated using the following homology arms:

5' Homology Arm:
GCCTCTGCACGCACATGCTTGCCTGTCAAACTCTACCACCCCGGCACGCTCTCTGTCTCC
3' Homology Arm:
AACTTACAAGAGGCCTTGTTCCGCTTCCCGGTATGGTAATTAGAAACTCATTAATGGGCG

Following induction of the λ Red recombinase, transformed cells were plated on LB agar supplemented with Chl, Kan, and Amp and incubated at 37°C. Positive clones were expanded and validated by Sanger sequencing. Next, markers were excised using Flp and Dre site-specific recombinases. Following induction, cells were plated on LB agar supplemented with Chloramphenicol. To confirm the loss of the selection cassettes, colonies were screened for sensitivity to Kanamycin and Ampicillin by parallel culture in LB broth containing 1) Chloramphenicol alone, 2) chloramphenicol and Kanamycin, and 3) Chloramphenicol, Kanamycin, and Ampicillin. Clones demonstrating growth exclusively in Chloramphenicol-supplemented media were subjected to PCR analysis. Successful recombination was identified by the presence of a 1089 bp recombination-specific band and the concomitant loss of Kan-specific (1039 bp) and Amp-specific (923 bp) internal sequences (**Fig. 2**). The final modified regions were validated by Sanger sequencing to confirm the presence of the expected residual FRT and Rox sites.

| Primer | Theoretical band (bp) | Identification Site | |
|---|---|---|---|
| GCAGGAGCCGCGCATATATACG | 627 | Wild-type Specific Site | Primers used to screen colonies for successful FRT-Kan-FRT and Rox-Amp-Rox selection cassettes into the two defined *ICP34.5* loci. |
| AAGAATATCATTAGCATGCACG | | | |
| ATAACGGTTTGGTTGATGCGAG | 1039 | Kan Resistance Gene-specific Site (Recombined) | |
| AAGAATATCATTAGCATGCACG | | | |
| AAGTGCTCATCATTGGAAAACG | 923 | Amp Resistance Gene-specific Site (Recombined) | |
| GGTGTTGTTACACCTGTGGCAG | | | |
| CCCAATGGCCGCGCGTCCCAGG | 2573/2314 | Full-length Site (Kan/Amp Segment) | |
| GGTGTTGTTACACCTGTGGCAG | | | |
| ATAACGGTTTGGTTGATGCGAG | 1039 | Kan Resistance Gene-specific Site | Primers used to screen colonies for successful dual *ICP34.5* deletion |
| AAGAATATCATTAGCATGCACG | | | |
| AAGTGCTCATCATTGGAAAACG | 923 | Amp Resistance Gene-specific Site | |
| GGTGTTGTTACACCTGTGGCAG | | | |
| GGTGTTGTTACACCTGTGGCAG | 1089 | Recombination-specific Site | |
| CCCAATGGCCGCGCGTCCCAGG | | | |

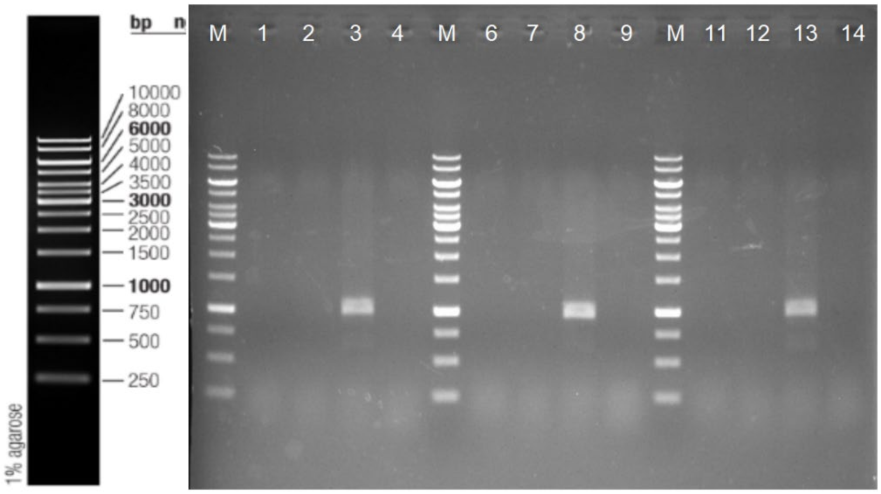


**Figure 2**: Colony PCR Gel Electrophoresis, *ICP34.5* deletion. Lanes 1, 6, and 11 represent kanamycin resistance specific site for three respective clones. Lanes 2, 7, and 12 represent ampicillin resistance specific site; lanes 3, 8, and 13 represent recombination-specific site. All three clones exhibited distinct recombination-specific bands corresponding to the predicted molecular size.

*ICP6::LacZ*: The *ICP6* (UL39) coding region was modified to insert a *LacZ-SV40pA* reporter construct using a two-step λ Red-mediated recombination protocol, per the HSV G47Δ design.

A ccdB-amp cassette was introduced into the coding region utilizing the temperature-sensitive helper plasmid pSC101-ccdA-gbaA. The targeting fragment was generated using the following homology arms:
5' Homology Arm:
CCGCTGGTGAGCCGGTCCGCTCGCCTTTACCGCATCCTGGGGGTTCTGGTGCACCTGCGG
3' Homology Arm:
ATCCGGACCCGGGAGGCCTCCTTTGAGGAGTGGCTGCGATCCAAGGAAGTGGCCCTGGAC

Recombination was induced, and cells were plated on LB agar supplemented with Chl, Amp, Tet, and L-arabinose at 30°C. Clones were screened via colony PCR using primers GGCGCTGGTCTTGCGGCCGATC and GCAGCTGGAGGTGTAGTAGTTG. Successful integration was identified by a 2325 bp recombination-specific band and the absence of the 903 bp wild-type band. Verified clones were cryopreserved and validated by Sanger sequencing.

The ccdB-amp cassette was subsequently replaced with a LacZ-SV40pA reporter construct. Following induction, cells were plated on LB agar containing Chloramphenicol and incubated at 37°C. Clones were validated by parallel PCR screening. Successful replacement was confirmed by the loss of the 1146 bp ccdB-amp internal fragment with primers: ATAACGGTTTGGTTGATGCGAG (Forward); AAGAATATCATTAGCATGCACG (Reverse) and the appearance of a 1392 bp recombination-specific band with primers: CAGTTCACCCGTGCACCGCTGG (Forward); GCAGCTGGAGGTGTAGTAGTTG (Reverse), **Fig. 3**. Sanger sequencing confirmed the integrity of the insertion, noting a synonymous C>T mutation in the LacZ insert.

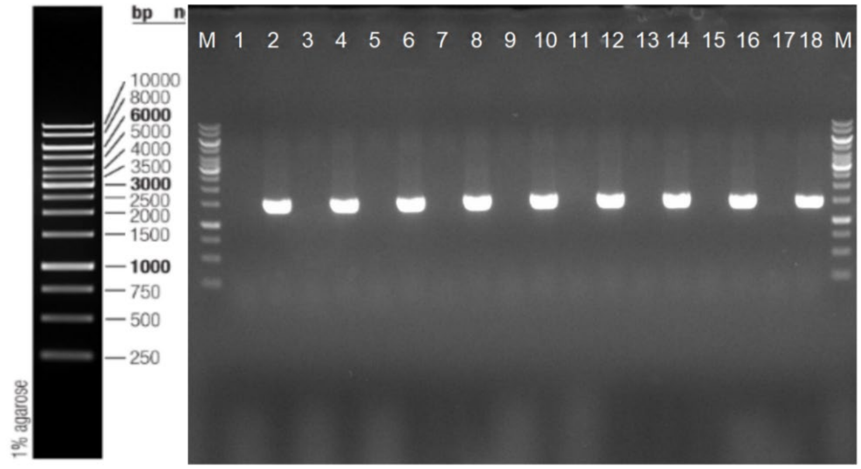

**Figure 3**: Colony PCR Gel Electrophoresis, *ICP6::LacZ*. Odd numbered lanes represent intermediate site with ccdB-amp selection cassette. Even lanes represent recombination-specific events. All 9 clones demonstrate successful recombination.

The verified BAC plasmids were extracted and transformed into E. coli DH10B. Transformants were selected on LB-Chl agar at 37°C. Final structural integrity was confirmed by PCR using primers GGCGCTGGTCTTGCGGCCGATC (Forward); GCAGCTGGAGGTGTAGTAGTTG (Reverse), yielding a predicted 4176 bp product (**Fig. 4**). Comprehensive Sanger sequencing confirmed 100% identity with the theoretical sequence for the modified regions.

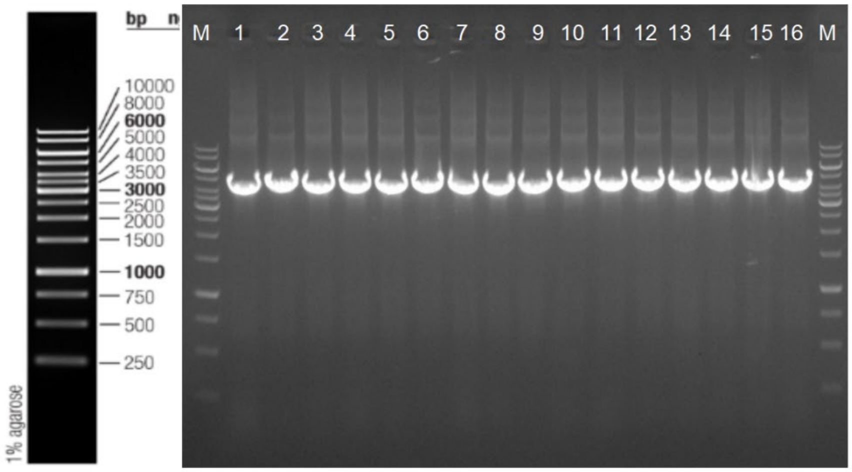


**Figure 4**: Colony PCR Gel Electrophoresis, *ICP6::LacZ*. 4176 bp band size encompassing the *ICP6::LacZ* modification is verified in the final DH10B transformed bacterial artificial chromosome (pBeloBAC11-HSV-1(F, ΔICP47_363bp, ΔICP34.5, ICP6::LacZ)

Functional validation of pBeloBAC11-HSV-1(F, ΔICP47_363bp, ΔICP34.5, ICP6::LacZ)

The validated BAC construct was propagated in *E. coli* and purified using the Qiagen Large-Construct Kit according to the manufacturer's instructions. Purified BAC DNA was transfected into Vero cells using the Lonza Nucleofector 4D X-unit (SF kit, pulse code DN-100). Following cytopathic effect (CPE) observation, the recombinant virus (designated HSV G47$\Delta^{Tk_null}$) was harvested and amplified according to standard protocols.[33] LacZ expression was confirmed via X-gal staining during plaque titration (**Fig. 5A**).[33]

Functional validation of the ICP34.5 and ICP47 deletions were performed in KNS42 cells infected at a multiplicity of infection (MOI) of 0.1 with HSV-1(F) (wild-type control), HSV G47Δ, or HSV G47$\Delta^{Tk_null}$. To validate the loss of ICP34.5 function, cell lysates were collected 24 hours post-infection and analyzed via Western blot for phosphorylated eIF2α. Functional ICP34.5 in wild-type HSV-1 prevents phosphorylation; therefore, successful deletion was defined by the induction of p-eIF2α in cells infected with HSV G47Δ and HSV G47$\Delta^{Tk_null}$ compared with its absence in wild-type infection (**Fig. 5B**). To validate the loss of ICP47-mediated immune evasion, infected KNS42 cells were harvested at 24 hours and stained with Spider β-gal (Dojindo) to identify infected cells, an anti-MHC Class I antibody (Enzo, ALX-805-711), and cell viability (Invitrogen, 65-0865-14). MHC Class I surface expression was quantified on viable, β-galactosidase-positive cells. Wild-type HSV-1 infection is characterized by the downregulation of surface MHC Class I. Consequently, functional deletion of ICP47 was determined by the retention of MHC Class I expression (comparable to uninfected controls) in cells infected with HSV G47Δ and HSV G47$\Delta^{Tk_null}$ compared to its reduction following wildtype infection (bulk viable population given negative β-gal), (**Fig. 5C**).

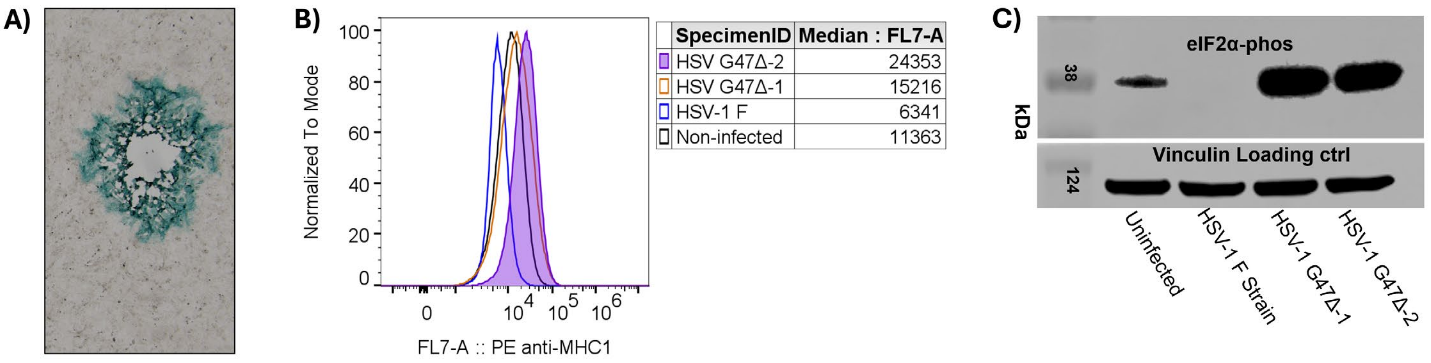


**Figure 5**: Functional validation of pBeloBAC11-HSV-1(F, ΔICP47_363bp, ΔICP34.5, ICP6::LacZ) modifications. **A)** Plaque assay demonstrating β-galactosidase expression (*ICP6::LacZ*) in the recombinant virus. **B)** Retention of MHC class 1 in infected KNS42 cells (*ΔICP47*). **C)** The recombinant HSV vector with dual *ICP34.5* deletion fails to suppress phosphorylation of eIF2α in infected KNS42, confirming functional dual locus deletions. HSV G47Δ-1 corresponds to native HSV G47Δ gifted by Dr. Jianfang ning. HSV G47Δ-2 corresponds to the recombinant HSV G47Δ derived from pBeloBAC11-HSV-1(F). HSV-1 F strain is the wildtype, unmodified virus purchased from ATCC.

Generation of pBeloBAC11-HSV-1(F,G47Δ,ΔICP34.5::rev(pBIRC5-ICP34.5-CBh-tTS))

The modified *ICP34.5* loci were further engineered to insert SV40pA:pBIRC5>ICP34.5:rBGpA - CBh>tTS:SV40pA into the deleted two-copy ICP34.5 loci of pBeloBAC11-HSV-1(F,G47Δ).

The 269-base pair *BIRC5* promoter sequence is:
CGCGTTCTTTGAAAGCAGTCGAGGGGGCGCTAGGTGTGGGCAGGGACGAGCTGGCGCGGCGTCGCTGG
GTGCACCGCGACCACGGGCAGAGCCACGCGGCGGGAGGACTACAACTCCCGGCACACCCCGCGCCGC
CCCGCCTCTACTCCCAGAAGGCCGCGGGGGGTGGACCGCCTAAGAGGGCGTGCGCTCCCGACATGCCC
CGCGGCGCGCCATTAACCGCCAGATTTGAATCGCGGGACCCGTTGGCAGAGGTGGCGGCGGCGGCA

Prior to insertion, the parental BAC glycerol stock was validated to confirm the integrity of the *ICP34.5* flanking regions. Cells were streaked on LB agar supplemented with Chloramphenicol (Chl) and incubated at 37°C. Single colonies were screened via colony PCR using the primers listed. Verified clones were expanded and cryopreserved.

| Primer sequence | Theoretical band (bp) | Identification loci |
|---|---|---|
| CCCAATGGCCGCGCGTCCCAGG | 1089 | homology arm |
| GGTGTTGTTACACCTGTGGCAG | | |
| GGTGTTGTTACACCTGTGGCAG | 1011 | homology arm |
| AAGAATATCATTAGCATGCACG | | |
| CAGTTCACCCGTGCACCGCTGG | 1392 | homology arm |
| GCAGCTGGAGGTGTAGTAGTTG | | |

A targeting fragment containing the reverse-oriented expression cassettes (SV40pA:pBIRC5>ICP34.5:rBGpACBh>tTS:SV40pA) linked to FRT-Kan-FRT and Rox-Amp-Rox selection markers was generated. The fragment utilized the following homology arms:
5' Homology Arm:
GCCTCTGCACGCACATGCTTGCCTGTCAAACTCTACCACCCCGGCACGCTCTCTGTCTCC
3' Homology Arm:
AACTTACAAGAGGCCTTGTTCCGCTTCCCGGTATGGTAATTAGAAACTCATTAATGGGCG

Transformed cells were selected on LB agar supplemented with Chloramphenicol, Kanamycin, and Ampicillin at 37°C. Clones were screened via colony PCR. Successful insertion was defined by the presence of a 6274 bp full-length band, a 3716/2512 bp *Kan*-specific band, and a 3745 bp *Amp*-specific band. A chosen clone was further validated by Sanger sequencing; approximately 20 bp within the GC-rich resistance gene loci could not be resolved due to technical limitations, but all other regions confirmed the correct sequence integration.

| Primer sequence | Theoretical band (bp) | Identification loci |
|---|---|---|
| GGTGTTGTTACACCTGTGGCAG | 6274 | Recombination-positive full-length locus |
| AAGAATATCATTAGCATGCACG | | |
| TCGAATTCCGAAGTTCCTATTCTCT | 3716/2512 | KanR-specific recombination-positive locus |
| CGGTCTAATCCTCAGGTGCA | | |
| TTCAAGGATCCTAACTTTAAATAATTGG | 3745 | AmpR-specific recombination-positive locus |
| CGGTCTAATCCTCAGGTGCA | | |

To generate the final construct, the antibiotic selection markers were excised via Flp and Dre recombinase activity. Following induction, cells were plated on LB agar containing Chloramphenicol only. Clones were screened for the loss of Kanamycin and Ampicillin resistance by parallel culture in selective media. Successful excision was confirmed by colony PCR, which demonstrated a shift in the full-length band size from 6274 bp to 5078 bp, corresponding to the loss of the selection cassettes (**Fig. 6**).

| Primer sequence | Theoretical band (bp) | Identification loci |
|---|---|---|
| TCGAATTCCGAAGTTCCTATTCTCT | 4192 | FRT-specific locus |
| AAGAATATCATTAGCATGCACG | | |
| TTCAAGGATCCTAACTTTAAATAATTGG | 4192 | Rox-specific locus |
| AAGAATATCATTAGCATGCACG | | |
| GGTGTTGTTACACCTGTGGCAG | 5078 | Recombination-positive full-length locus of Intermediate BAC |
| AAGAATATCATTAGCATGCACG | | |

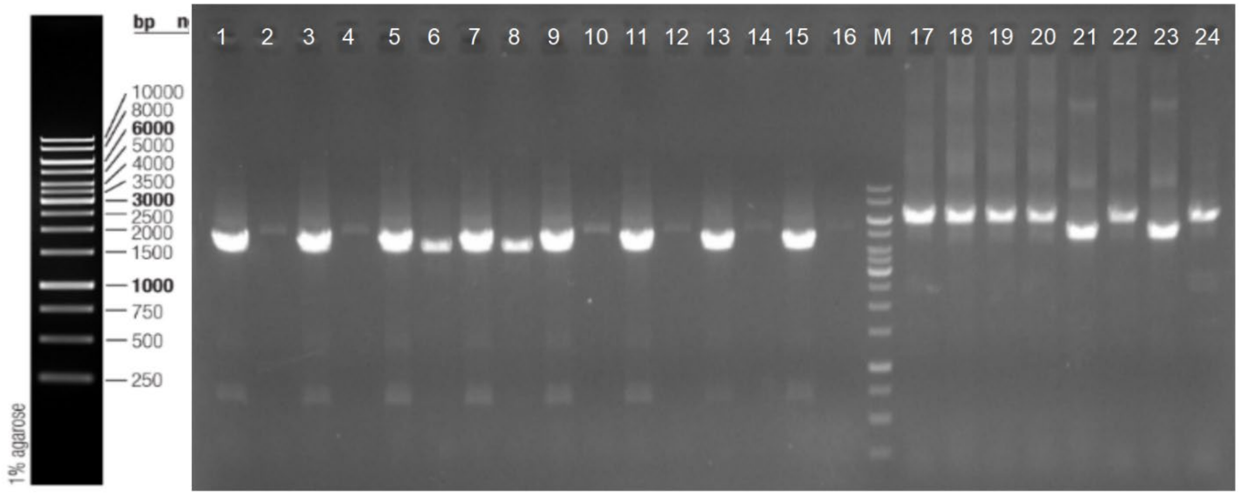


**Figure 6**: Colony PCR Gel Electrophoresis, *ΔICP34.5::rev(pBIRC5-ICP34.5-CBh-tTS)*. Lanes 1, 2, and 17 correspond to a single clone (and repeated for a total of eight clones), with the first band corresponding to FRT-specific locus, second band to Rox-specific locus, and third band to the recombination-positive full length locus. Clones 3 (lanes 5, 6, and 21) and clone 4 (lanes 7, 8, and 23) showed bands of the expected sizes for successful recombination.

The sequence of the final modified loci was analyzed via Sanger sequencing. While high GC-content in the cassette resulted in a ~120 bp unresolvable region, the flanking and internal coding sequences were confirmed to match the theoretical design (100% identity). To definitively validate the global integrity of the vector, the final clone was subjected to Circular Sequencing, which confirmed the correct architecture of the complete BAC plasmid.

<u>Functional validation of pBeloBAC11-HSV-1(F,G47Δ,ΔICP34.5::rev(pBIRC5-ICP34.5-CBh-tTS))</u>

The validated BAC construct was propagated in *E. coli* and purified using the Qiagen Large-Construct Kit according to the manufacturer’s instructions. Purified BAC DNA was transfected into Vero cells using the Lonza Nucleofector 4D X-unit (SF kit, pulse code DN-100). Following cytopathic effect (CPE) observation, the recombinant virus was harvested and amplified according to standard protocols.[33]

To validate functional restoration of *ICP34.5* activity under the *BIRC5* promoter, S462-TY cells were infected at a multiplicity of infection (MOI) of 0.1 with HSV-1(F) (wild-type control), HSV G47Δ, or HSV G47$\Delta^{Tk_null}$, ins(*BIRC5>ICP34.5-CBh-tTS*). Cell lysates from two biological replicate experiments were collected 24 hours post-infection and analyzed via Western blot for phosphorylated eIF2α, normalized to total eIF2α. As shown (**Fig. 7**), both HSV-1 F strain wildtype virus and pBeloBAC11-HSV-1(F,G47Δ,ΔICP34.5::rev(pBIRC5-ICP34.5-CBh-tTS) successfully repressed p-eIF2α:eIF2α, while induction is retained following HSV G47Δ infection.

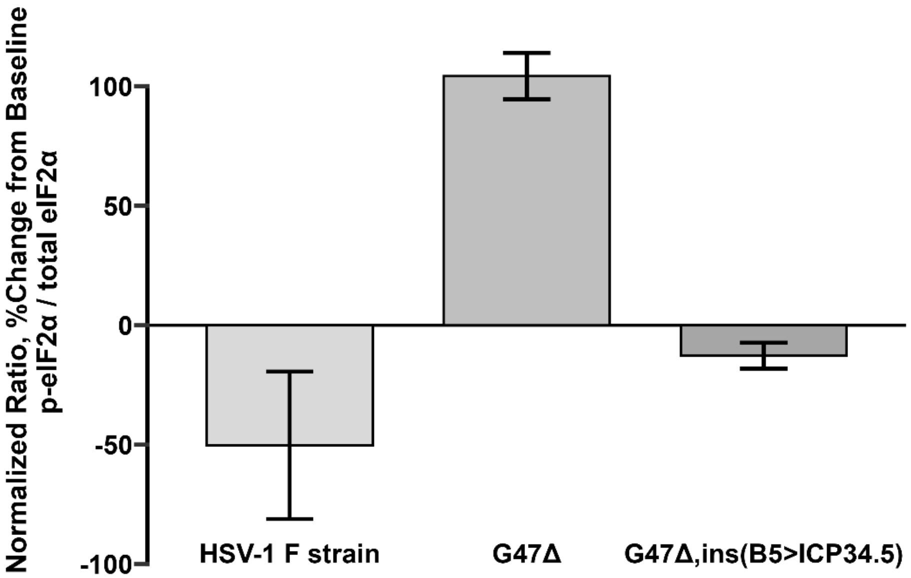


**Figure 7:** Validation of conditional *ICP34.5* restoration in infected S462-TY cells. The ratio of p-eIF2α:eIF2α is expressed as a percentage compared to uninfected control.

Validation of UL23 restoration in HSV G47$\Delta_2$ and or HSV G47$\Delta_2$, ins(*BIRC5>ICP34.5-CBh-tTS*)

In the original pBeloBAC11-HSV-1(F), the viral *UL23* (*TK*) gene is interrupted with the bacterial gene elements required for maintenance in *E. coli* (referred to as BAC elements). The 5’ BAC element contains a repeated sequence corresponding to the initial portion of the *UL23-2* sequence immediately downstream of the BAC elements (**Figure 8a**). This permits homologous recombination when the plasmid is transfected into eukaryotic cells for viral particle production, to excise the unneeded BAC elements while restoring the *UL23* open reading frame. To validate UL23 open-reading-frame restoration, S462-TY cells were seeded onto 6-well plates. Wells were infected with HSV G47Δ, HSV G47$\Delta_2$, and HSV G47$\Delta_2$,ins(*BIRC5>ICP34.5-CBh-tTS*), respectively, at 0.05 MOI for 48 hours. After 48 hours, genomic DNA was purified from the pellets. Flanking *UL23* primers (UL23_Primer1: *TCAGTTAGCCTCCCCCATCT* and UL23_Primer2: *TTCCGAGACAATCGCGAACA*) were used to evaluate for *UL23* restoration via PCR and gel electrophoresis in cells infected by the corresponding viruses, with an expected band size of 853 base pairs. Primers for GAPDH were used as a control (Forward: *CTTTGTCAAGCTCATTTCCTGGT*; Reverse: *AGATGAAAAGAGTTGTCAGGGC*), with an expected band size of 120 base pairs. Results revealed intact *UL23* in all viruses (**Figure 8b**).

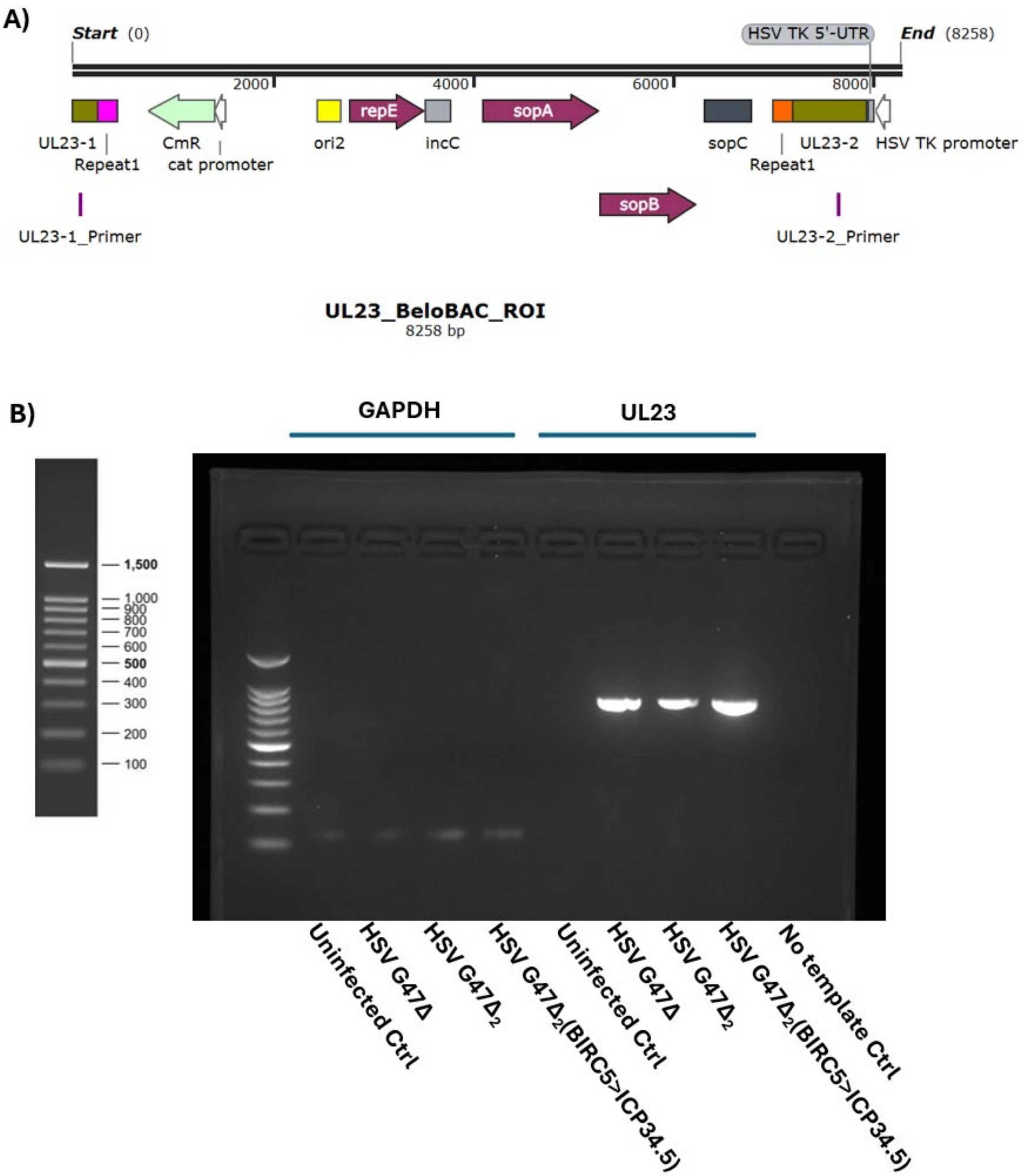


**Figure 8**: **A)** pBeloBAC11-HSV-1(F) design. The *UL23* region of interest (ROI) contains the *UL23* gene (*UL23-1:* and *UL23-2:*) interrupted by the BeloBAC backbone. Repeat regions (Repeat1, magenta and orange) permit homologous recombination excision upon transfection into eukaryotic host cells to restore the *UL23* open reading frame. **B)** Flanking *UL23* primers (UL23_Primer1: *TCAGTTAGCCTCCCCCATCT* and UL23_Primer2: *TTCCGAGACAATCGCGAACA*) were used to evaluate for *UL23* restoration in cells infected by the corresponding viruses. The expected *UL23* band size (853 bp) is verified in HSV G47Δ (positive control), HSV G47$\Delta_2$, and HSV G47$\Delta_2$, ins(*BIRC5>ICP34.5-CBh-tTS*). *GAPDH* PCR was used for control (expected band size = 120 bp).
